# BioPrediction-PPI: Simplifying the Prediction of Protein-Protein actions through Artificial Intelligence

**DOI:** 10.1101/2025.11.16.688401

**Authors:** Bruno R. Florentino, Robson P. Bonidia, Ulisses Rocha, André C. P. L. F. de Carvalho

## Abstract

Proteins are essential in biological processes, primarily through their interactions with other molecules, including proteins. These interactions are crucial for cellular functions and maintaining life. Predicting Protein-Protein Interactions (PPIs) is very important, although challenging, for understanding cellular functions and diseases. This paper presents BioPrediction-PPI, a new end-to-end Machine Learning (ML) framework for PPI prediction, automating the entire process from feature extraction to interpretability, with no manual intervention required. BioPrediction-PPI stands as one of the few end-to-end models that do not rely on deep learning, enabling the automated use of trained models on new data. It offers interpretability graphs for creating auditable models and has been evaluated through comparative experiments with 30 previous studies, using 15 datasets. Additionally, the interpretability graphs offer valuable insights for model evaluation and experimental design, facilitating informed decision-making. BioPrediction-PPI demonstrates competitive performance across multiple datasets, even without the use of deep learning, and is a transparent, white-box model that can be easily used by biologists and practitioners without a background in computer science. The proposed framework has the potential to accelerate research in biology and related fields by making AI-driven tools more accessible.

## 1 Introduction

Proteins are the most versatile molecules, typically performing their functions through interactions with other molecules, primarily other proteins [44], in protein-protein interactions (PPIs). These interactions are fundamental to cellular pathways and the maintenance of life [9]. Among the functions that these interactions can perform are DNA synthesis [9, 51, 44], gene transcription and translation [9, 51, 44], drug signaling [9], immune response [9, 44], enzymatic catalysis [9], post-transcriptional modification of proteins [51], viral infection mechanisms [46], and cell proliferation [44], to name a few examples.

Throughout history, various viruses have caused deadly diseases. A recent and notable example is SARS-CoV-2, the causative agent of COVID-19, which triggered a global pandemic [8]. The SARS-CoV-2 infection process involves binding viral cells to a receptor in human cells through a spike protein in a PPI, highlighting the importance of understanding this type of interaction to understand the infection and its consequences [17]. In this sense, studies on the relationships between the SARS-CoV-2 proteins and human proteins revealed that these interactions are related to biological processes such as translation, transcription, and regulation, in addition to helping to identify pharmaceutical targets [17, 1].

The same issue can be extended to other infectious viruses that also utilize human proteins as hosts [46]. This includes diseases caused by herpes viruses, HIV, hepatitis, dengue, and Zika, since understanding the interaction between the virus and the host is crucial for comprehending the infectious mechanism, the infections themselves, and potential pathways for drug development [15].

An additional example highlighting the importance of understanding PPIs is their application in the reverse vaccinology approach. In this context, information derived from interaction networks is often used to identify the most suitable target proteins for vaccine development [16]. Studies in this field have shown that analyzing these networks enables the identification of highly connected proteins, which typically play essential roles in maintaining metabolism and pathogen pathogenicity [23, 5].

Furthermore, investigating these interactions provides insights into the role of these proteins within their respective pathways, enabling a detailed analysis of the interactions involved, as well as the submolecular context and functions they perform [2, 5, 22, 36]. This approach helps determine which proteins should proceed to subsequent stages of vaccine development, such as selecting peptides with immunogenic potential, while also contributing to an in-depth study of promising candidates. This strategy has been applied to various pathogenic organisms, including Plasmodium falciparum [2], Chlamydia trachomatis [5], XDR Salmonella typhi H58 [23], and Aeromonas hydrophila [22].

However, studies of this nature often rely on the STRING interaction database for data retrieval, which may limit the analysis to proteins and interactions that have been previously cataloged on this platform. So, in this context, the discovery of new PPIs plays a crucial role in the process of developing therapies and drugs. This is because identifying new PPI targets opens up opportunities for novel therapeutic approaches [10, 51, 47].

Even so, experimental methods are often expensive and time-consuming to perform [10, 9, 44, 20], creating a growing demand for computational techniques to predict these interactions [9], such as Machine Learning (ML). Nevertheless, working with biological sequences using computational approaches presents several challenges. A fundamental obstacle to applying ML techniques to protein sequences lies in the data, which is inherently categorical and unstructured [7].

Despite the widespread availability of numerous ML libraries and platforms accessible and free to all users, many individuals lack the necessary skills to confidently begin their journey into project development using ML [14]. In this context, a discourse emerges regarding the democratization of Artificial Intelligence (AI), encompassing various dimensions such as democratization in model development [32]. This democratization has the potential to accelerate technological innovations [41]. Such endeavors can include the provision of models through open-source channels and the utilization of automated pipelines [32, 38, 41].

Automated solutions for predicting PPIs are highly attractive to researchers in the field of biological sciences, especially those with little or no knowledge of programming, which makes it difficult to develop predictive models independently. In this context, the low requirements for knowledge in data science encourage the adoption of ML models in areas of study where computational methods are not yet widely used.

In this regard, robust models capable of handling different dataset configurations are an important aspect. This is because, in many cases, it is not possible to build a dataset with tens of thousands or millions of interactions directly related to the problem. Therefore, a robust model can be applied under unfavorable data conditions while maintaining satisfactory performance. Another aspect that can contribute to the usefulness of a model is its high generalization ability. In other words, this means that, even when trained with a specific dataset, the model can predict a wide range of interactions that are not directly related to the original problem.

Moreover, another important challenge in the use of ML models is the lack of interpretability, a critical factor, especially in the medical and biological fields, where decisions based on model results can directly impact individuals’ health [4]. The perception of models as “black boxebecause data needs to be transformed into meaningful features that ML algorithms can use later to make inferences, such as classifying between interacting andr more traditional ML models, rather than more complex approaches like Deep Learning (DL), can be a beneficial strategy, as these models are generally simpler and offer greater interpretability. Furthermore, having a deep understanding of a model’s performance is crucial for evaluating the reliability of its predictions and determining which results are truly useful for decision-making [3, 4].

With this in mind, we propose BioPrediction-PPI, an end-to-end ML framework for predicting PPIs. As a fully automated framework, users must input data, with all steps, from feature extraction to model interpretability, being handled automatically. Considering this, the research questions (RQs) of this article are as follows:

**RQ1:** Is it feasible to develop an end-to-end ML framework for predicting PPIs that eliminates the need for intervention by ML experts while remaining competitive in performance with models developed by specialized professionals?

**RQ2:** Is the developed framework, being automated, capable of maintaining performance and robustness under conditions of unbalanced data and varying dataset sizes?

**RQ3:** Is the developed framework capable of generating models with high generalization, trained for one context, but able to perform well in other contexts?

**RQ4:** Is it possible to develop an interpretable end-to-end model that generates reports clarifying the model’s decision patterns, which can assist in the decision-making process for users?

Therefore, this study proposes an automated framework for predicting PPIs, where four aspects will be validated: (1) its performance as an automated framework, (2) robustness across datasets with different configurations, (3) generalization to unseen contexts, and (4) interpretability of the patterns generated by the models. In this sense, BioPrediction-PPI^1^ ^2^ can significantly impact the development of ML by non-experts, as it was designed to be both automated and interpretable. Thus, it contributes to the advancement of research related to metabolism and provides a deeper understanding of the pathways involved in diseases.

## 2 Materials and Methods

### 2.1 Related Works

Numerous options have been validated in different contexts in state-of-the-art studies for predicting PPIs, using only the primary structure of protein characterization. A notable example among white-box models is LigthGBM-PPI [9]. This stands out for its ability to extract features from protein sequences, employing strategies such as pseudo amino acid composition, autocorrelation descriptor, and local descriptor [9]. Subsequently, it selects the best features using an elastic net and, finally, trains the model using LigthGBM, which is based on decision trees [9].

Similarly to the previous model, we have GTB-PPI [51], which extracts a feature vector using techniques such as pseudo amino acid composition (PseAAC), pseudo-position-specific scoring matrix (PsePSSM), and autocorrelation descriptor (AD) [51]. Subsequently, to reduce dimensionality, L1-regularized logistic regression (L1-RLR) is used, and then the best tree-based model is selected for the model training [51].

Furthermore, most current models in the literature utilize DL models. One of the leading models is PIPR, capable of predicting PPIs solely using the primary structure of each protein, with a deep residual recurrent convolutional neural network in the Siamese architecture that considers the influence of both molecules on the problem [10]. In the same context, Yang et al. [46], which adopts a Siamese architecture of Convolutional Neural Network (CNN) to extract features from each sequence and a Multi-Layer Perceptron (MLP)-based model for decision-making [46].

Another DL model is DeepFE-PPI [47], which utilizes a proprietary technique called Res2vec, based on Word2vec, to represent biological sequences. Additionally, it uses a Deep Neural Network (DNN) to classify the tasks into interaction or non-interaction categories [47]. In contrast to previous models, the model by Yang et al. [44], besides using DL for classification and extracting structural information from sequences, also extracts topological features from them [44]. LSTM-PHV [40] is a DL prediction tool that uses the Long Short-Term Memory (LSTM) model and feature extraction through word2vec embedding. Unlike most existing models, this tool applied a pre-trained model in the context of human and pathogenic interactions hosted on a web server that allows new predictions for any user [40].

Furthermore, the TUnA model [24] is a DL model that relies exclusively on structural information for interaction prediction, explicitly focusing on cross-species predictions. This means the model is trained for one species and aims to predict PPIs for others. TUnA uses the ESM-2, a pre-trained protein language model, which extracts 640 * *n* features from each protein, where *n* is the number of amino acids in the sequence, making it a highly dimensional method [24]. Finally, it employs the Spectral-normalized Neural Gaussian Process (SNGP) to perform the predictive task.

Table 1 highlights the key features of the studies used in the performance comparison. Although some studies may exhibit variations, as outlined in the results tables, they fundamentally share the same core characteristics. This analysis highlights the availability of the models, as some provide only experimental descriptions in the article, others have links that are no longer functioning, and others have folders with the code. Additionally, it indicates whether the tool has documentation explaining how to use the codes, with ’no’ for those that have no documentation teaching how to run the tool, ’reproduction’ for documentation that teaches how to execute with the datasets described in the work, and ’new exp.’ for documentation that teaches how to apply it to new data.

**Table 1:** The key features of the state-of-the-art studies. First, the ‘Available’ and ‘Documentation’ categories assess whether a solution is accessible and well-documented to support new users. The documentation is classified into three levels: “no” (not available), “reproduction” (allows only the replication of original experiments), and “new exp” (new experiment). (supports experiments with new data). Next, the ‘End-to-End Fitting’ category identifies studies that claim to develop fully automated tools capable of handling all modeling steps for new data, and the ’End-to-End Prediction’ category identifies tools that allow the use of the fitted models to predict new interactions. The ‘Deep Learning’ column highlights tools that use deep learning models for predictions. The ‘Additional Analyses’ category indicates studies that provide outputs beyond pairwise interaction predictions, such as interpretability results. Finally, ‘Studies Compared’ and ‘Datasets Compared’ measure the extent of validation in each study, reflecting the number of comparative analyses conducted.

| Tool | Available | Documentation | End-to-End Fitting | End-to-End Prediction | Deep Learning | Additional Analyses | Studies Compared | Datasets Compared |
| --- | --- | --- | --- | --- | --- | --- | --- | --- |
| SVM-AC [18] | no | no | no | no | no | no | 2 | 3 |
| EELM-PCA [48] | no | no | no | no | no | no | 5 | 2 |
| SVM-MCD [49] | no | no | no | no | no | no | 8 | 2 |
| SAE [37] | no | no | no | no | no | no | 8 | 11 |
| DNN-PPI [25] | no | no | no | no | yes | no | 2 | 7 |
| PIPR [10] | git-hub | new exp.<sup>a</sup> | yes<sup>b</sup> | no | yes | no | 11 | 4 |
| DeepPPI [13] | no | no | no | no | yes | no | 15 | 10 |
| RF-LPQ [42] | no | no | no | no | no | no | 3 | 1 |
| LD+KNN [45] | no | no | no | no | no | no | 0 | 1 |
| LightGBM-PPI [9] | git-hub | no | no | no | no | no | 13 | 6 |
| MIMI+NMBAC+RF [12] | online folder | no | no | no | no | no | 16 | 8 |
| LRA+RF [50] | no | no | no | no | no | no | 8 | 2 |
| GTB-PPI [51] | git-hub | no | no | no | yes | no | 10 | 7 |
| DeepFE-PPI [47] | git-hub | reproduction | no | no | yes | no | 22 | 7 |
| DPPI [19] | git-hub | new exp.<sup>a</sup> | no | no | yes | no | 11 | 6 |
| DeepTrio [20] | web and git | new exp.<sup>a</sup> | no | no | yes | yes<sup>d</sup> | 6 | 5 |
| LGB+BBA [52] | git-hub | no | no | no | no | no | 10 | 2 |
| DeepCF-PPI [39] | git-hub | no | no | no | yes | no | 20 | 10 |
| Huang et al. [33] | no | no | no | no | no | no | 12 | 9 |
| Liu et al. [26] | online folder | offline | no | no | no | no | 0 | 1 |
| DCT+WSRC [21] | no | no | no | no | no | no | 16 | 3 |
| Signature products [28] | no | no | no | no | no | no | 3 | 4 |
| HKNN [29] | no | no | no | no | no | no | 5 | 2 |
| Ensemble of HKNN [30] | no | no | no | no | no | no | 4 | 2 |
| Yang et al. [46] | git-hub | no | yes<sup>b</sup> | no | yes | no | 0 | 8 |
| Emvirus [43] | git-hub | no | no | no | no | no | 1 | 8 |
| LSTM-PHV [40] | web server | new exp.<sup>a</sup> | yes<sup>b</sup> | yes<sup>c</sup> | yes | no | 2 | 2 |
| D-SCRIPT [35] | git-hub | new exp.<sup>a</sup> | yes<sup>b</sup> | no | yes | no | 1 | 6 |
| Topsy-Turvy [34] | git-hub | new exp.<sup>a</sup> | yes<sup>b</sup> | no | yes | no | 3 | 6 |
| TUNA [24] | git-hub | new exp.<sup>a</sup> | yes<sup>b</sup> | no | yes | no | 7 | 7 |
| **BioPrediction-PPI** | **git-hub** | **new exp.<sup>a</sup>** | **yes<sup>b</sup>** | **yes<sup>c</sup>** | **no** | **yes<sup>d</sup>** | **30** | **16** |

<sup>a</sup> documented studies that teach how to apply them to new datasets for model fitting. <sup>b</sup> only fully automated tools for model fitting.
<sup>c</sup> Only tools that allow using trained models on new data without coding. <sup>d</sup> Only studies that provide complementary reports to the user.

The upcoming features concern models that claim to be end-to-end (i.e., automated tools capable of running the entire pipeline). These features will be divided into two parts, ’end-to-end fitting’, for studies capable of adjusting a predictive model in an automated way, only with the input of data for training, and ’end-to-end prediction’, for those tools that, after adjusting the predictive model, have an automated way of using this model in new interactions.

Next, we have the column for additional categories, where we can see which tools, in addition to predicting interactions, provide some additional analysis output that could guide a potential user. It is also mentioned whether the methodology includes DL techniques. Finally, the table also indicates the number of expert-developed models included in the comparison and the datasets utilized for validation.

We observed that a larger proportion of the studies, approximately 47% (14 of 30 studies), lack code available to new users. This issue arises because the authors did not share their code, and the referenced sites are no longer online. In the other cases where the code is available, it can be found on GitHub, other online repositories, or hosted on dedicated web servers. Additionally, we saw that almost half of the studies that make their code available lack documentation, which deters non-programmer users.

Only six of the 30 studies analyzed are end-to-end, all of which are based on DL. BioPrediction-PPI is the only end-to-end framework that does not use DL. Furthermore, the framework developed in this work is the only one that provides code in the documentation for reapplying the trained model to predict new pairs, unlike other models that claim to be end-to-end, where automation ends once the model is trained and performance is estimated. Thus, no code is available for using these models to predict new protein pairs. In parallel, we have the server with the pre-trained model, LSTM-PHV [40], which can be used to predict new interactions. However, it only allows the use of a specific pre-trained model, without adapting it to the particular problem at hand.

Moreover, the only other tools that offer additional analyses are DeepTrio [20] and BioPrediction-PPI. The former, DeepTrio [20], provides a figure that highlights which amino acids in the protein sequences were most important for the final decision. On the other hand, BioPrediction-PPI includes performance graphs by threshold for the user and feature importance graphs aimed at interpretability.

This scenario poses a barrier to the democratization of AI, as most studies remain out of reach for users who lack expertise in programming and ML. The barriers include a lack of code availability, poor documentation, and reliance on non-interpretable techniques. Together, these factors make it difficult for users to use PPI prediction tools in real-world applications, which would otherwise accelerate studies related to protein interactions. As our proposal is an end-to-end ML framework intended for use in multiple contexts by users, this validation process with 30 studies and 15 datasets serves as a central pillar to report the robustness of the proposed framework.

### 2.2 Methodology

BioPrediction-PPI predicts PPIs with four key focuses: automation, robustness, generalization, and interpretability. The first experiment will evaluate the performance of the end-to-end framework on three benchmarks, comparing its prediction capability with existing models and assessing the performance obtained against other models developed by experts. Next, the robustness of the model will be analyzed by validating its performance on different datasets of the same subproblem, with unbalanced data and varying sizes, allowing for an evaluation of the consistency of the results.

Next, the generalization of the models will be evaluated for prediction in new contexts, through a model based on human interactions and the prediction of interactions from five other species. Finally, the interpretability of the model will be investigated in a case study, assessing its outputs, the ability to explain the learned patterns, and provide biological insights (Figure 1).

**Figure 1:**
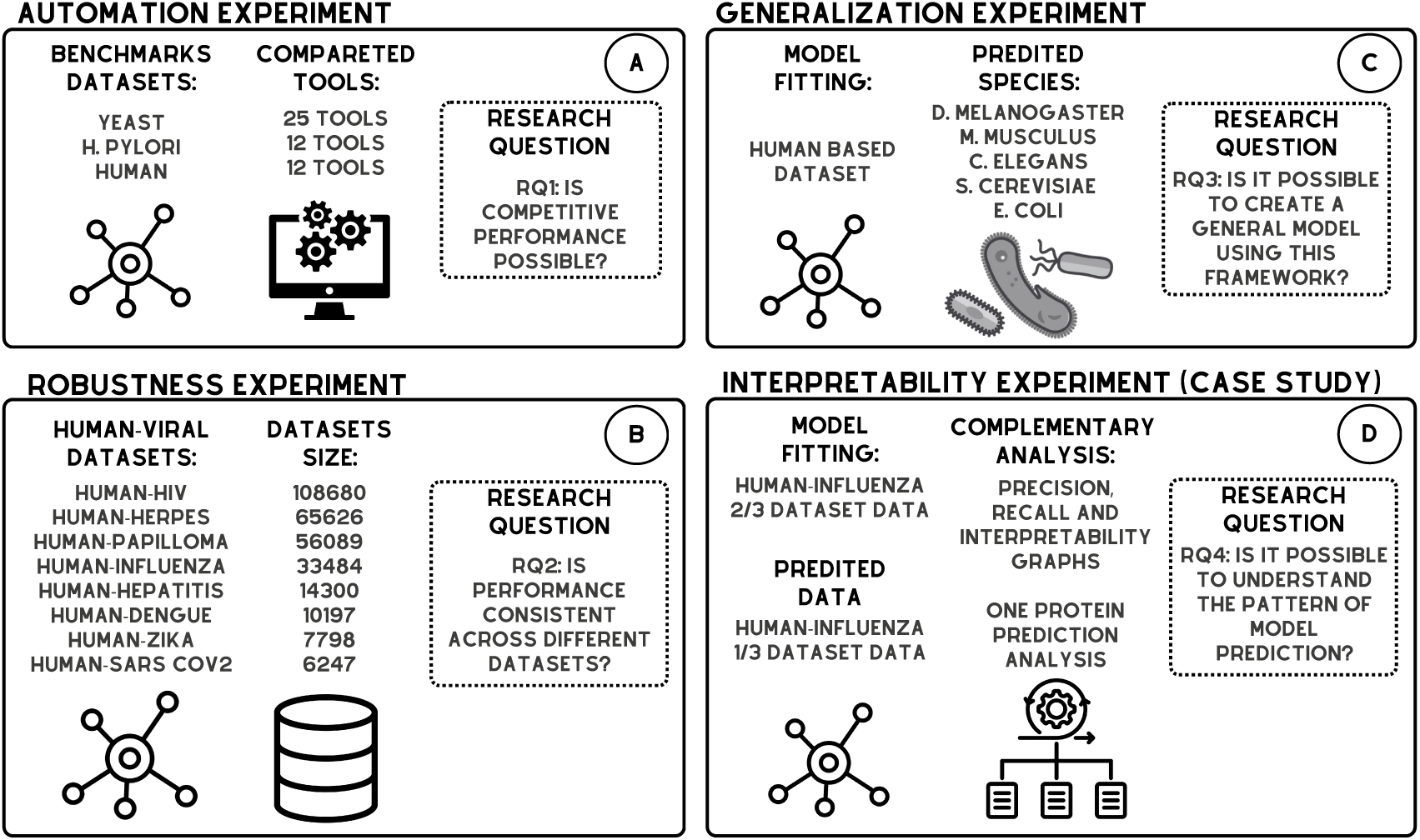
Experiments design: In the automation experiment (A), the framework’s performance is assessed using the benchmarks of yeast, H. pylori, and human, comparing each performance with 24, 12, and 12 other studies, respectively. Next, the robustness experiment (B) is evaluated using unbalanced data (1:10 ratio) and datasets from the same subproblem, with sizes ranging from approximately 6,000 to 100,000 interactions. In the generalization experiment (C), the capacity of the models is also measured by training with human interaction data and using the model to predict interactions from five other species. Finally, in the interpretability experiment (D), the human-influenza interaction dataset is used, with 2/3 of the data allocated for model training and the remaining 1/3 for representing candidate interactions. Subsequently, the model’s complementary outputs regarding the learned patterns is analyzed.

#### 2.2.1 Experimental Setup

Accuracy is the fraction of correct predictions relative to the total number of predictions made. Precision, on the other hand, measures the proportion of correct positive predictions relative to the total number of positive predictions made. Recall evaluates the model’s ability to identify all positive instances. Additionally, specificity checks the model’s ability to identify negative instances correctly. The F1-score, in turn, is the harmonic mean between precision and recall, providing a balance between them.

In addition to these metrics, the MCC is a metric that evaluates the overall quality of a binary classifier, taking into account all four prediction categories: True Positives, True Negatives, False Positives, and False Negatives. AUROC measures the area under the ROC (Receiver Operating Characteristic) curve, which displays the true positive rate relative to the false positive rate. Finally, AUPRC measures the area under the Precision-Recall curve, which relates precision to recall. This metric is particularly useful in imbalanced datasets, as it provides a more robust evaluation of the model.

In the comparison process, BioPrediction-PPI is considered non-competitive with another tool if, for individual metric comparisons, more metrics show superior performance for the other tool than those where BioPrediction-PPI’s performance is comparable. This comparison is determined through a one-tailed t-test. The test takes into account the mean, standard deviation, and sample size, with a significance level (alpha) of 0.05. The null hypothesis assumes that the mean performance of the evaluated tool is less than or equal to BioPrediction-PPI’s mean performance. The alternative hypothesis, however, posits that the evaluated tool’s mean performance is greater than BioPrediction-PPI’s. On the other hand, BioPrediction-PPI is deemed competitive when the majority of the framework’s metrics are equal to or greater than those of the compared tool.

In cases where the author has not reported the standard deviation for the metrics, the non-parametric Mann-Whitney U test will be performed unilaterally, with a significance level (alpha) of 0.05, comparing the means of all metrics simultaneously. The null hypothesis posits that the mean performance of the evaluated tool is not statistically superior to that of BioPrediction-PPI (i.e., BioPrediction-PPI is competitive). In contrast, the alternative hypothesis asserts that the mean performance of the evaluated tool is superior to that of BioPrediction-PPI.

### 2.3 Validation Experiments

The BioPrediction-PPI’s performance was validated by comparing it with others workflows cited in Table 1 using various datasets described in Table 2. Four experiments were conducted, adhering to methodologies described in the experimental setup subsection, which are compatible with the methodologies of the four studies used for comparison (1).

**Table 2:** Datasets used in the experiments. For each dataset, we provide the number of positive interactions, negative interactions, and total interactions, along with information on which experiment the dataset belongs to.

| Dataset | Total interactions | Positive interactions | Negative interactions | Experiment |
| --- | --- | --- | --- | --- |
| Yeast | 11188 | 5594 | 5594 | One |
| H. pylori | 2916 | 1458 | 1458 |  |
| Human | 8102 | 3882 | 4220 |  |
| HIV | 108680 | 9880 | 98800 | Two |
| Herpes | 65626 | 5966 | 59660 |  |
| Papilloma | 56089 | 5099 | 50990 |  |
| Influenza | 33484 | 3044 | 30440 |  |
| Hepatitis | 14300 | 1300 | 13000 |  |
| Dengue | 10197 | 927 | 9270 |  |
| Zika | 7798 | 709 | 7090 |  |
| Sars Cov2 | 6247 | 568 | 5680 |  |
| Human cross | 474517 | 43138 | 431379 | Three |
| D. melanogaster | 55000 | 5000 | 50000 |  |
| E. coli | 55000 | 5000 | 50000 |  |
| C. elegans | 55000 | 5000 | 50000 |  |
| S. cerevisiae | 55000 | 5000 | 50000 |  |
| M. musculus | 20000 | 2000 | 22000 |  |
| Influenza model | 19949 | 1686 | 18263 | Four |
| Influenza candidates | 13535 | 1358 | 12177 |  |
| P03496 test set | 45 | 22 | 23 |  |
| P03496 unlabeled | 16190 | - | - |  |

#### 2.3.1 Automation Experiment Methodology

For the development and testing of computational approaches, benchmark datasets enable quality comparisons with other similar tools [31]. Therefore, the first experiment utilized benchmark datasets [10, 9] to evaluate the performance of BioPrediction-PPI in predicting unknown interactions of an organism, using previously known interactions as input, and comparing its performance with other tools developed by experts.

This evaluation is crucial because, to develop a tool that can be used by non-experts in data science, it is not enough for the tool to be automated; it must also be as efficient as existing tools. Therefore, in this experiment, the performance of BioPrediction-PPI will be evaluated against other PPI prediction studies that utilize only the primary structure of proteins to adjust predictive models.

The **first dataset** is a yeast-based dataset and will be used for comparison with 24 tools that describe your performance in 7 studies [10, 9, 51, 47, 20, 52, 39], evaluating six distinct metrics (Acc, Prec, Rec, Spec, F1, MCC).

In the **second dataset**, representing H. pylori PPIs, BioPrediction-PPI is compared with 12 tools in four different metrics (Acc, Rec, Prec, MCC) [9, 51, 47]. Finally, the **third dataset** will be conducted with human PPIs compared to 12 tools in four different metrics (Acc, Rec, Prec, MCC) [13] [47]. Additionally, the first two datasets are balanced, while the third is slightly imbalanced, as described in Table 2.

#### 2.3.2 Robustness Experiment Methodology

The main goal of the **second experiment** is to evaluate and understand the robustness of the framework. In this experiment, human-viral interactions will be used, with eight datasets corresponding to eight different species, to assess the versatility and consistency of the framework in a specific subtopic. See Table 2 for the species and dataset sizes. In other words, the goal is to determine whether BioPrediction-PPI can predict PPIs for all species or only for some of them. Additionally, each dataset has distinct sizes, allowing for the evaluation of performance evolution as the number of known interactions increases, with all datasets having a 1:10 ratio. Therefore, the ability to handle various viral species, with different dataset sizes, under imbalanced data conditions will be assessed.

In this experiment, to compose the reference performances for these datasets, BioPrediction-PPI is compared with three tools from the literature: two described in articles reporting their performances through cross-validation [46] [43], and a third, the LSTM-PHV web server, which specializes in predicting interactions between human and pathogen proteins [40]. The LSTM-PHV server was used to predict known interactions, and its performance was subsequently evaluated. It is essential to note that the LSTM-PHV server only supports proteins with fewer than 1000 amino acids, resulting in a reduction of up to 20% in the total interactions included for evaluation. For each tool, six different metrics (Acc, Prec, Rec, Spec, F1, MCC) were calculated across eight different datasets, each corresponding to a different viral species.

#### 2.3.3 Generalization Experiment Methodology

Next, in **third experiment**, the generalization of the models trained by BioPrediction-PPI will be evaluated. Specifically, we will assess how well the model, once trained on a dataset from one species, can predict interactions in other species. This is a particularly challenging task in interaction prediction because models often learn extensively about the specific context in which they are trained. However, they may fail to identify general patterns and instead only understand the training context, leading to poor performance when applied outside that initial context [24]. Therefore, in a **cross-species experiment**, the generalization of BioPrediction-PPI was tested.

To address this, a state-of-the-art benchmark was used to evaluate cross-species performance, which involves training a model on human interactions and then predicting interactions for E. coli, M. musculus, D. melanogaster, C. elegans, and S. cerevisiae (See Table 2). Performance comparisons were made with six tools, five of which have their performance reported in TUnA [24], and the sixth is the performance of the LSTM-PHV Web server [40], with AUPR and AUROC metrics being considered.

In this experiment, cross-validation was not performed because the author of TUnA [24] provided specific training and test sets. Consequently, BioPrediction-PPI was adapted to skip cross-validation. Instead, the provided training set was further divided into training and validation subsets. Subsequently, the final performance was calculated using the set of tests provided. This approach ensured that all tools were consistently evaluated on the same data.

#### 2.3.4 Interpretability Experiment Methodology - Case Study

After validating the performance under different conditions, a case study will be conducted using one of the datasets to assess the interpretability of the framework. This approach is useful not only for gaining biological insights from the learned patterns but also for selecting the most appropriate set of predicted interactions to be considered as positive for future experiments. When model performance is similar, the one that provides more information about the learned patterns may be more valuable, as it aids in planning new experiments by guiding the selection of the most relevant interactions to test and validate.

For such, the complementary graphs of the framework will be explored. Specifically, we opted for the Influenza dataset, which comprises 33,484 interactions, including 3,044 positive and 30,440 negative interactions. Initially, we divided the interactions into two subgroups with a 6/10 and 4/10 ratio: one for model construction and the other to illustrate potential user interaction candidates, while maintaining the proportions of positive and negative classes.

In the candidate subset, interactions can occur only between protein pairs that are present in the training subset, eliminating the possibility of proteins appearing exclusively in the candidate set. This creates a dataset that simulates the scenario of a user aiming to predict new connections in the interaction network that have not yet been discovered.

Then, the predictive model will be trained using BioPrediction-PPI, and the candidate interactions will be predicted using the first fold of the model. This will allow for a comparison between the performance estimated during the modeling process and the performance obtained for this 4/10 of the data, evaluating whether the model maintains performance close to the expected, as measured using the standard threshold of 0.5.

Additionally, the graphs generated with the modeling data will be compared to their corresponding versions created from the candidate interaction data. These graphs examine the model’s patterns to enhance understanding of its functionality, as well as the estimated performance using a 0.5 threshold. The goal is to assess the feasibility of using the output graphs from BioPrediction-PPI to estimate the model’s behavior when predicting new interactions. Since the candidate interactions have known labels, this analysis will validate whether the graphs truly reflect the expected performance in new prediction scenarios.

Finally, a specific analysis will be conducted on the results for the protein P03496 from the Influenza virus, which stands out due to having a large number of positive interactions in the candidate interaction set. This will allow for the construction of the distribution of predicted scores for this protein. From this distribution, the consistency of the predictions for this protein will be discussed in relation to the general patterns of the model.

Additionally, a dataset of unlabeled interactions will be constructed, including all unlabeled interactions between P03496 and human proteins present in the dataset. These interactions will then be predicted using the first fold of the trained model. The coherence of the predictions made for this unlabeled set will be discussed, considering the limitations imposed by the absence of labels, while evaluating whether the predictions follow similar patterns to the labeled interactions and whether they could indicate potential new relevant interactions. Table 2 contains more information about the datasets used in this experiment.

### 2.4 BioPrediction-PPI: Workflow

Our proposal adopts an automated procedure for creating end-to-end ML models aimed at predicting interactions, along with generating reports designed to investigate specific characteristics of the model. To initiate the model-building process, it is essential to provide the necessary data, comprising two main elements: a file listing the known interactions and a dictionary containing the sequences of all relevant proteins. The list of interactions is presented in a tabular format, containing three columns: the names of the involved proteins and an indication of the interaction type. The protein dictionary, in turn, is a FASTA-formatted document containing all the primary protein sequences related to the problem.

Internally, the initial process involves subdividing the known interactions into three distinct sets: training, validation, and testing, with proportions of 70%, 10%, and 20%, respectively. The model’s performance is evaluated at the end based on the predictions made with the test set. Additionally, the cross-validation technique is applied with five folds, which implies evaluating the performance in five different subdivisions to further validate the model on the data. Next, the process advances to the feature extraction module, where information is obtained to describe the protein sequences. These features are classified into two main types: structural (see Figure 2a) and topological (see Figure 2b). Structural features are those derived directly from the primary sequences of each protein and include aspects such as amino acid frequencies, Shannon entropy, and statistical measures of the Fast Fourier Transform (FFT) [6]. On the other hand, topological features are extracted from the interaction network, which is present exclusively in the training set. These include interaction counts and other graph metrics, such as centrality and betweenness.

**Figure 2:**
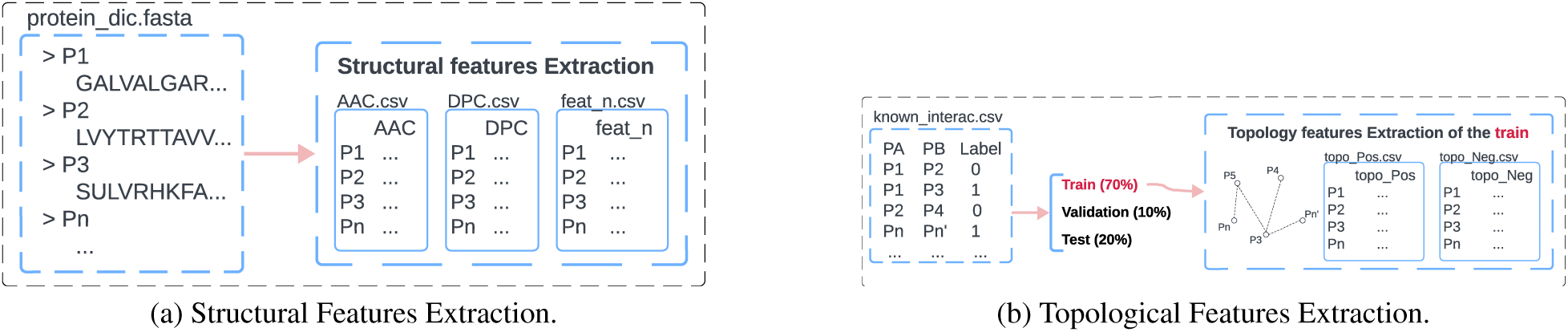
Feature extraction scheme for protein characterization.

The next step involves constructing the datasets for the modeling phase by concatenating the protein features with the interaction table. This results in a table where each row corresponds to the features of two proteins, and the label indicates the relationship or interaction associated with that pair. Each feature descriptor generates a distinct dataset, as indicated at the left of Figure 3. In the subsequent step, partial models are trained for each feature descriptor, aiming to reduce the problem’s dimensionality and consequently increase efficiency in the final execution. The training and validation sets are utilized in this process, with the former for training and the latter for evaluating performance.

**Figure 3:**
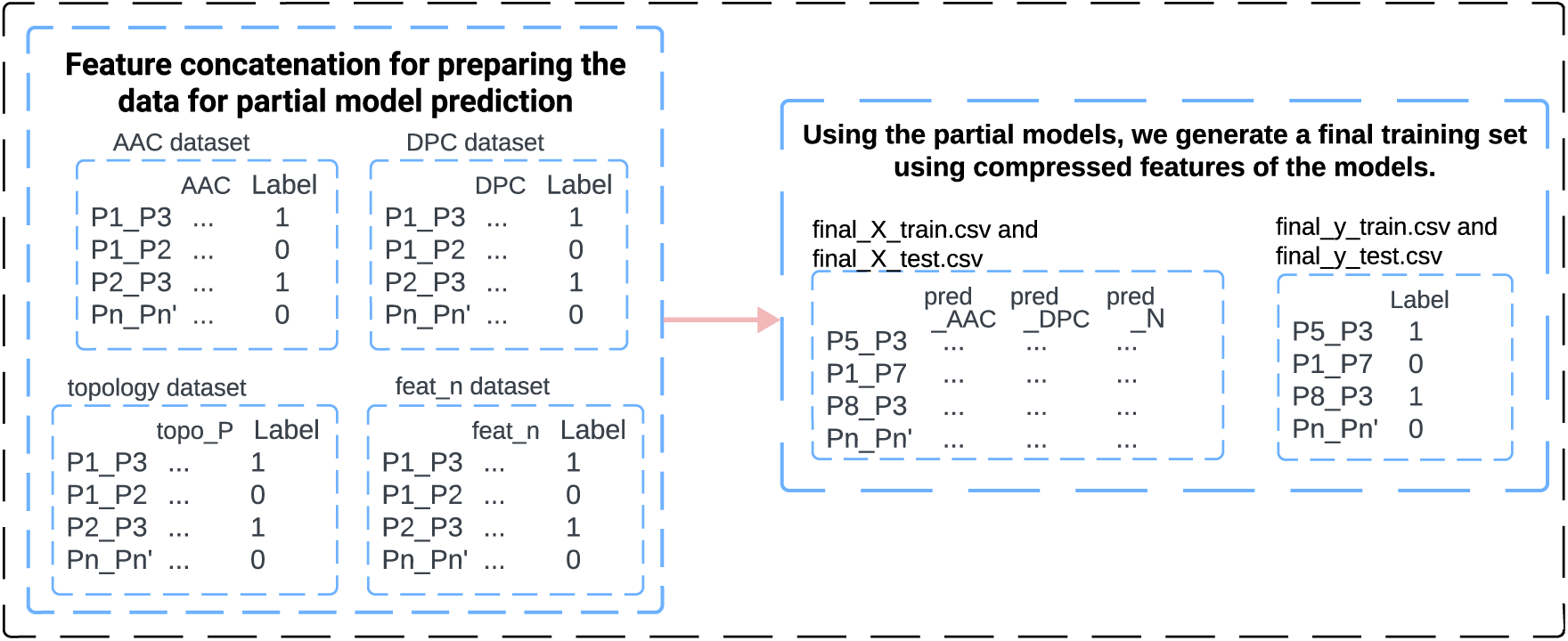
Illustration of the process: feature concatenation to create partial datasets, fitting partial models, and combining partial decisions into a final dataset.

After constructing these partial models, the probability of belonging to the interaction class is used as the new compressed feature. For example, when using amino acid composition (AAC) to build a partial model, initially comprising 20 columns, the predicted probability from this partial model is utilized as the new compressed AAC. This process is repeated for all feature descriptors, culminating in the formation of a final dataset containing all compressed features for definitive training and testing. In this phase, the validation set is employed as the training set. In contrast, the test set is used to evaluate the performance of the final model, thus preventing information leakage by using a previously unobserved test set. The workflow of partial models is also schematized in Figure 3.

The final training and test sets are used to construct the model that definitively determines the class of each interaction pair; this dynamic is shown in Figure 4. The intermediate and final models are constructed using decision tree-based algorithms, including Random Forest, CatBoost, and XGBoost.

**Figure 4:**
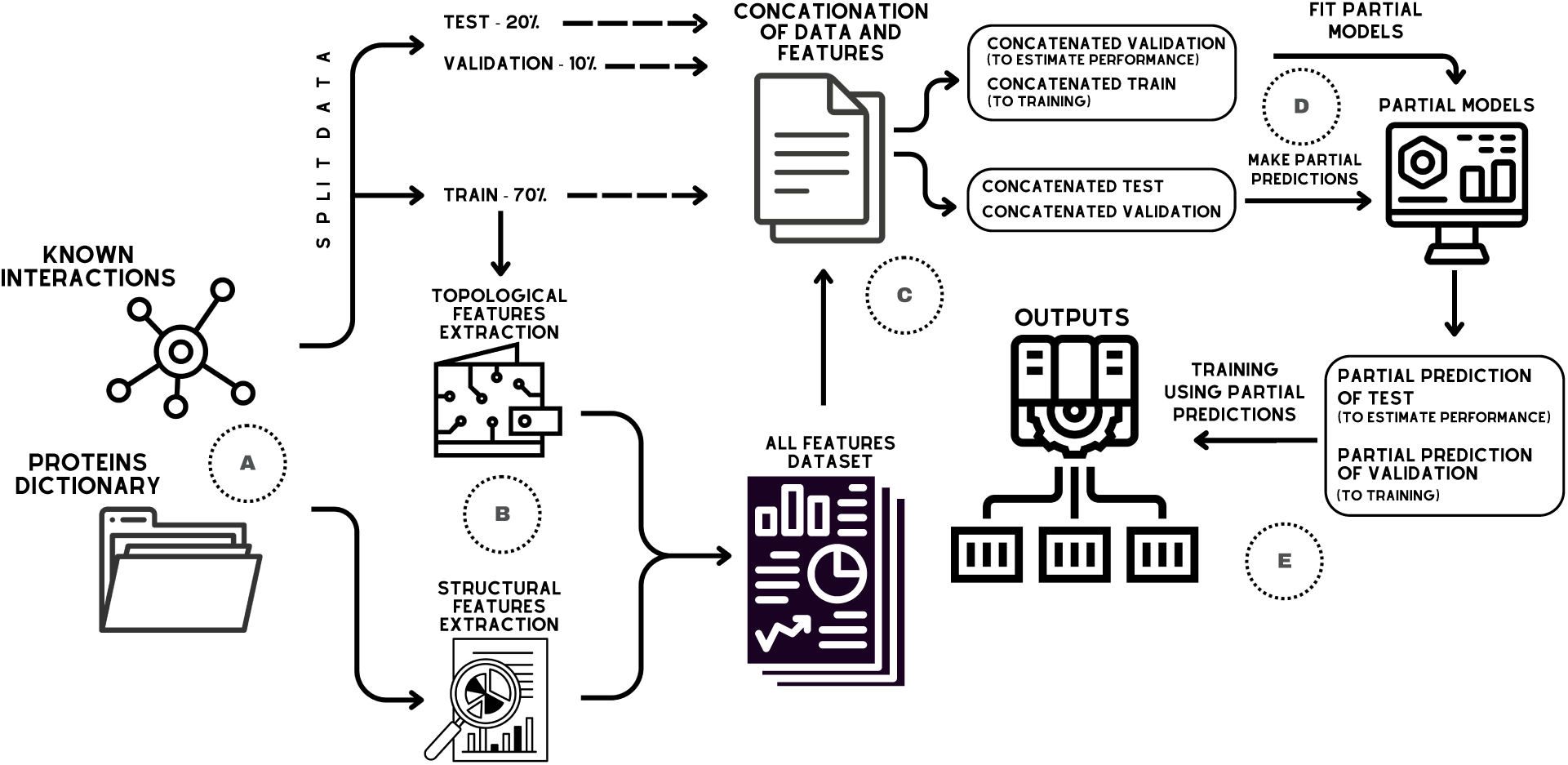
Workflow of BioPrediction-PPI: In A, inputs are entered; in B, feature extractions occur, where the training data is used to extract topological features and the protein dictionary to extract structural features; in C, training, validation, and test datasets are concatenated with the extracted features; in D, training and validation data for each feature descriptor are employed to fit partial models, afterwards, validation and testing are predicted by the partial models. Finally, in E, decisions from partial models are combined to fit a final model, resulting in the delivery of the model for PPI prediction and other relevant outcomes.

#### 2.4.1 BioPrediction-PPI Reports

With the model prepared, an interpretability report is generated using the SHAP Values library [27]. This interpretability report employs various graphs to facilitate the understanding of the impact each feature has on the model’s decision-making process. These visual representations are designed to identify patterns and analyze the relationship between the magnitude of a feature and its corresponding class, highlighting both the most relevant contributions to classification and the influence of the feature value distributions across different classes.

Additionally, a usability report is generated to provide the user with information about the model’s metrics and properties. All these graphs are generated based on the test set, the same dataset used to evaluate the model’s performance. This ensures that the same behavior is expected when predicting new interactions, making these graphs useful for assisting in the selection of which newly predicted pairs should be considered in the planning of future experiments.

For such, graphs are generated to help understand the fitted model, as each predicted interaction pair is assigned a score ranging from 0 to 1. Conventionally, interactions with a score equal to or greater than 0.5 are considered positive, and machine learning model metrics are calculated based on this symmetrical threshold. However, the distribution of these scores may reveal relevant patterns, enabling a more in-depth analysis of the model’s behavior.

The first available graph is the **distribution of predicted scores**. In this graph, the distribution of predicted scores is plotted separately for each class, considering all protein pairs. In other words, all positive pairs from the test subset are first selected, and their predicted score distribution is constructed. The same procedure is then applied to the negative class.

In an ideal model, all positive pairs should be distributed with scores above 0.5, while negative pairs should fall below this threshold. However, in real models, some positive pairs may receive scores lower than this threshold, indicating misclassification. Analyzing the overlap region between the two class distributions can provide insights into how the model assigns scores to actual classes and highlight the regions where misclassification is more likely to occur.

Next, we present two graphs related to the analysis of the positive class precision in the fitted model. The precision of the positive class corresponds to the percentage of correct classifications among those that the model predicts as positive. For example, a precision of 80% for a decision threshold of 0.5 (considering the range between 1.0 and 0.5) means that, out of every 100 predictions with a score above 0.5, approximately 80 belong to the positive class.

The first graph used to assess the model’s precision is the **precision by threshold interval**. In this graph, the precision of the positive class is calculated for different score intervals: (1.0–0.9), (0.9–0.8), (0.8–0.7), (0.7–0.6), and (0.6–0.5). The goal of this analysis is to determine whether there is a specific interval where precision is significantly higher than the overall average, enabling more reliable predictions to be made. For example, suppose a fitted model has an overall precision of 60% for the conventional range (1.0–0.5). However, when considering only predictions in the (1.0–0.9) interval, the precision may reach 95%. In this case, even if the overall precision is not very high, there exists a region where the model correctly classifies almost all predictions. This means that, depending on the user’s objective, it is possible to leverage part of the model’s predictions by focusing only on pairs classified within this high-confidence interval.

Additionally, we perform an analysis of **precision by threshold**, which calculates cumulative precision from a score of 1.0 down to a given threshold. Unlike the previous graph, which calculates intervals without overlap, this graph accumulates precision values, starting at 1.0 and progressively including predictions down to the chosen threshold. This approach estimates the precision obtained when testing all interactions from the highest score down to a specific point.

This graph considers the number of predictions in each interval. For example, if the first interval has a precision of 90% and the next one 70%, the graph will show the overall performance by weighting the number of predictions in each score range. While the first graph helps identify the most precise regions, this one assists in defining the optimal cutoff point, that is, the threshold up to which predictions maintain an acceptable level of precision for the user.

In parallel, there is also a graph related to the **sensitivity of the positive class** in the fitted model. In this context, sensitivity represents the percentage of total positive interactions that are correctly classified as belonging to the positive class. For example, a sensitivity of 80% for a decision threshold of 0.5 (considering the range between 1.0 and 0.5) means that if there are 100 positive interactions in total, 80 of them will have scores above 0.5; in other words, they will be classified as positive.

In this context, to explore the model’s sensitivity, we use the **sensitivity by threshold graph**, which calculates cumulative sensitivity from a score of 1.0 down to a given threshold. In other words, this graph displays the total percentage of positive interactions captured up to a specific cutoff point. The purpose of this graph is to provide a measure of how comprehensively the model captures positive interactions within a given range. Unlike precision graphs, which typically focus on the range from 1.0 to 0.5, sensitivity is analyzed across the entire spectrum, from 1.0 to 0.0. This allows for identifying the necessary cutoff threshold to achieve a desired minimum sensitivity.

For example, if the goal is to reach 80% sensitivity, it may be necessary to consider all interactions classified as positive (scores between 1.0 and 0.5), as well as a fraction of interactions originally predicted as negative (e.g., between 0.5 and 0.3). Thus, this graph helps determine the extent to which expanding the set of tested interactions is beneficial, balancing coverage and precision.

## 3 Results

### 3.1 Automation Experiment Results

The focus of the first experiment is to evaluate the performance of our end-to-end framework in comparison with other methods using three benchmark datasets. As previously mentioned, BioPrediction-PPI’s performance will be compared metric by metric against other studies. For this comparison, a hypothesis test will be applied, where the null hypothesis states that the performance of the metric in the compared study is equal to or lower than that of BioPrediction-PPI. If at least half of the analyzed metrics are equal to or lower than the framework, it will be considered competitive; otherwise, it will be classified as non-competitive.

In the first dataset using the yeast dataset, we observe that out of the 25 compared studies, BioPrediction-PPI is considered non-competitive in only one case. The complete results of dataset one are summarized in Table 3. The competitive exception applies to PIPR [10], as this tool exhibits superior performance in four of the six evaluated metrics. Although BioPrediction-PPI, given the proposed methodology, is classified as non-competitive, the difference in accuracy between this model and PIPR [10] does not exceed 2%, as our framework achieved an accuracy of 95.16 *±* 0.25, while the PIPR study [10] reported an accuracy of 97.09 *±* 0.24.

**Table 3:** Performance in automation experiment, in dataset one (yeast dataset), measuring the Accuracy (Acc), precision (Prec), recall (Rec), specificity (Spec), F1, and Matthews Correlation Coefficient (MCC). In bold are the performances where there is statistical evidence of performance superior to BioPrediction-PPI.

| Study | Acc | Prec | Rec | Spec | F1 | MCC | Ref |
| --- | --- | --- | --- | --- | --- | --- | --- |
| SVM-AC | 87.35 ± 1.38 | 87.82 ± 4.84 | 87.30 ± 5.23 | 87.41 ± 6.33 | 87.34 ± 1.33 | 75.09 ± 2.51 | [10] |
| EELM-PCA | 86.99 ± 0.29 | 87.59 ± 0.32 | 86.15 ± 0.43 | N/A | 86.86 ± 0.37 | 77.36 ± 0.44 | |
| SVM-MCD | 91.36 ± 0.4 | 91.94 ± 0.69 | 90.67 ± 0.77 | N/A | 91.3 ± 0.73 | 84.21 ± 0.66 | |
| MLP | 94.43 ± 0.3 | 96.65 ± 0.59 | **92.06 ± 0.36** | N/A | 94.3 ± 0.45 | 88.97 ± 0.62 | |
| RF-LPQ | 93.92 ± 0.36 | 96.45 ± 0.45 | 91.10 ± 0.31 | N/A | 93.7 ± 0.37 | 88.56 ± 0.63 | |
| SAE | 67.17 ± 0.62 | 66.90 ± 1.42 | 68.06 ± 2.50 | 66.30 ± 2.27 | 67.44 ± 1.08 | 34.39 ± 1.25 | |
| DNN-PPI | 76.61 ± 0.51 | 75.1 ± 0.66 | 79.63 ± 1.34 | 73.59 ± 1.28 | 77.29 ± 0.66 | 53.32 ± 1.05 | |
| SRGRU | 93.77 ± 0.84 | 94.60 ± 0.64 | **92.85 ± 1.58** | 94.69 ± 0.81 | 93.71 ± 0.85 | 87.56 ± 1.67 | |
| SCNN | 95.03 ± 0.47 | 95.51 ± 0.77 | **94.51 ± 1.27** | 95.55 ± 0.77 | **95.00 ± 0.50** | 90.08 ± 0.93 | |
| PIPR | **97.09 ± 0.24** | 97.00 ± 0.65 | **97.17 ± 0.44** | 97.00 ± 0.67 | **97.09 ± 0.23** | **94.17 ± 0.48** | |
| DeepPPI | 94.43 ± 0.30 | 96.65 ± 0.59 | 92.06 ± 0.36 | N/A | N/A | 88.97 ± 0.62 | [9] |
| MCD | 91.36 ± 0.36 | 91.94 ± 0.62 | 90.67 ± 0.69 | N/A | N/A | 84.21 ± 0.59 | |
| LD+KNN | 86.15 ± 1.17 | 90.24 ± 1.34 | 81.03 ± 1.74 | N/A | N/A | N/A | |
| LightGBM-PPI | 95.07 ± 0.51 | 97.82 ± 0.48 | **92.21 ± 0.61** | N/A | N/A | 90.30 ± 1.01 | |
| MLD+RF | 94.72 ± 0.43 | 98.91 ± 0.55 | **94.34 ± 0.49** | N/A | N/A | 85.99 ± 0.89 | [51] |
| MIMI+NMBAC+RF | 95.14 ± 1.80 | 97.10 ± 2.10 | **92.67 ± 0.50** | N/A | N/A | 90.10 ± 0.92 | |
| LRA+RF | 94.43 ± 0.30 | 97.10 ± 2.10 | 91.22 ± 1.60 | N/A | N/A | 88.96 ± 0.026 | |
| GTB-PPI | 95.15 ± 0.25 | 97.97 ± 0.60 | **92.21 ± 0.36** | N/A | N/A | 90.45 ± 0.53 | |
| DeepFE-PPI | 94.78 ± 0.61 | 96.45 ± 0.87 | **92.99 ± 0.66** | N/A | N/A | 89.62 ± 1.23 | [47] |
| DPPI | **95.55** | 96.68 | **92.24** | N/A | 94.41 | N/A | [20] |
| DeepDuo | 94.14 ± 0.30 | 96.37 ± 1.43 | 91.74 ± 1.26 | 96.51 ± 1.41 | 93.98 ± 0.27 | 88.40 ± 0.67 | |
| DeepTrio | 94.78 ± 0.28 | 97.18 ± 0.28 | 92.20 ± 0.49 | 97.33 ± 0.30 | 89.67 ± 0.55 | **94.63 ± 0.29** | |
| LGB + BBA | 94.39 | 92.26 | **96.36** | N/A | N/A | N/A | [52] |
| DeepCF-PPI | 95.6 ± 0.57 | 93.4 ± 0.69 | **97.81 ± 0.54** | N/A | N/A | **91.4 ± 1.13** | [39] |
| BioPrediction-PPI | 95.16 ± 0.25 | **98.38 ± 0.40** | 91.31 ± 0.25 | **98.68 ± 0.24** | 94.71 ± 0.29 | 90.24 ± 0.50 | |

Additionally, the developed framework demonstrates competitive performance against all evaluated white-box models and several black-box models, including DeepPPI, GTB-PPI, DeepTrio, and DeepCF-PPI. When compared with other studies, BioPrediction-PPI showed equal or superior performance in at least four of the six evaluated metrics, demonstrating competitiveness with 96% (23 of 24) of the tested studies.

In general, considering the 25 studies, a total of 113 individual metrics were compared. In 91 of these metrics, BioPrediction-PPI demonstrated equal or superior performance, corresponding to outstanding performance in 82% of the evaluated metrics. In other words, the other models only exhibit superior metric values in isolated cases. Among the 22 individual metrics in which BioPrediction-PPI was not competitive, 13 (or 59%) are related to recall, indicating that this is the area where the model performed the worst compared to existing studies. On the other hand, for the precision metric, none of the evaluated models demonstrated statistically significant superior performance.

In the second experiment, we used a benchmark dataset based on *H. pylori* to evaluate the performance of our proposal against 12 studies, as shown in Table 4. BioPrediction-PPI was competitive with all of them, except for one tool, LGB+BBA [52]. In other words, the only study that, after applying the hypothesis test, we found does not have performance equal to or inferior to BioPrediction-PPI. More specifically, in the second dataset, the model was 2% more accurate and 3% more precise compared to BioPrediction-PPI. However, in the first dataset, the model demonstrated performance equal to or inferior to BioPrediction-PPI, being 1% less accurate and 6% less precise.

**Table 4:** Performance in automation experiment, in dataset two (H. pylori dataset), measuring the Accuracy (Acc), precision (Prec), recall (Rec), and Matthews Correlation Coefficient (MCC). In bold are the performances where there is statistical evidence of performance superior to BioPrediction-PPI.

| Study | Acc | Rec | Prec | MCC | Ref |
| --- | --- | --- | --- | --- | --- |
| PCA-EELM | 87.50 | 88.95 | 86.15 | 78.13 | [9] |
| MCD | 84.91 | 83.24 | 86.12 | 74.40 | – |
| MMI+NMBAC | 87.59 | 86.81 | 88.23 | 75.24 | – |
| DCT+WSRC | 86.74 | 86.43 | **97.01** | 76.99 | – |
| Signature products | 83.40 | 79.90 | 85.70 | N/A | – |
| HKNN | 84.00 | 86.00 | 84.00 | N/A | – |
| Ensemble of HKNN | 86.60 | 86.70 | 85.00 | N/A | – |
| LightGBM-PPI | 89.03 | 89.99 | 88.36 | 78.14 | – |
| WSR | 83.70 | 79.00 | 87.00 | N/A | [51] |
| DeepPPI | 86.23 | 89.44 | 84.32 | 72.63 | – |
| GTB-PPI | 90.47 ± 0.84 | 91.15 ± 1.42 | 89.99 ± 2.06 | 81.00 ± 1.63 | – |
| DeepFE-PPI | 88.36 | 89.19 | 88.20 | N/A | [47] |
| LGB+BBA | **97.89** | **99.21** | **96.55** | N/A | [52] |
| BioPrediction-PPI | 94.05 ± 0.18 | 97.24 ± 0.24 | 93.24 ± 0.12 | **86.68 ± 0.46** | – |

**Table 5:** Performance in automation experiment, in dataset three (human dataset), measuring the Accuracy (Acc), precision (Prec), recall (Rec), and Matthews Correlation Coefficient (MCC). Performances were taken from DeepFE-PPI [47]. In bold are the performances where there is statistical evidence of performance superior to BioPrediction-PPI.

| Study | Acc | Rec | Prec | MCC |
| --- | --- | --- | --- | --- |
| DeepFE-PPI | **98.71 ± 0.30** | **98.54 ± 0.55** | **98.77 ± 0.53** | **97.43 ± 0.61** |
| Ding, Tang & Guo (2016) | 97.56 | 96.57 | 98.30 | 95.13 |
| Ding, Tang & Guo (2016) | 96.08 | 95.05 | 96.97 | 92.17 |
| Ding, Tang & Guo (2016) | 95.59 | 94.06 | 96.94 | 91.21 |
| DeepPPI - Du et al. (2017) | **98.14** | 96.95 | **99.13** | **96.29** |
| Huang et al. (2015) | 96.30 | 92.64 | **99.59** | 92.82 |
| Liu et al. (2015) | 96.40 | 94.20 | N/A | 92.8 |
| Liu et al. (2015) | 95.70 | **97.6** | N/A | 91.8 |
| Liu et al. (2015) | 90.70 | 89.7 | N/A | 81.3 |
| Liu et al. (2015) | 95.50 | 94.0 | N/A | 91.4 |
| Liu et al. (2015) | 95.10 | 93.3 | N/A | 91.0 |
| Liu et al. (2015) | 89.30 | 94.0 | N/A | 79.2 |
| BioPrediction-PPI | 97.56 ± 0.06 | 97.21 ± 0.08 | 97.69 ± 0.15 | 95.21 ± 0.11 |

For the third dataset, the objective was to evaluate the effectiveness of BioPrediction-PPI in predicting a human PPI benchmark. Among the 12 studies analyzed, BioPrediction-PPI was competitive with 10 of them, surpassed only by DeepFE-PPI [47] and DeepPPI [13], both black-box models. Nevertheless, our framework demonstrated competitive performance with these models in the yeast dataset, suggesting that the tools are competitive. Specifically, in the yeast dataset, BioPrediction-PPI achieved approximately 1% higher accuracy, whereas in this human dataset, these tools were about 1% more accurate.

After conducting the experiments with three benchmarks, the results indicate that BioPrediction-PPI demonstrates performance equal to or superior to most of the compared models. When there is a performance difference, the variation in accuracy does not exceed 2%, suggesting that the use of the proposed end-to-end framework does not result in models inferior to those of previous approaches.

### 3.2 Robustness Experiment Results

In the robustness experiment, we utilized eight datasets comprising interactions between human proteins and viruses. In this analysis, BioPrediction-PPI demonstrated competitiveness across all compared datasets (see Table 6), indicating robustness, as it remained competitive with the performance of other existing studies, regardless of the viral species.

**Table 6:**
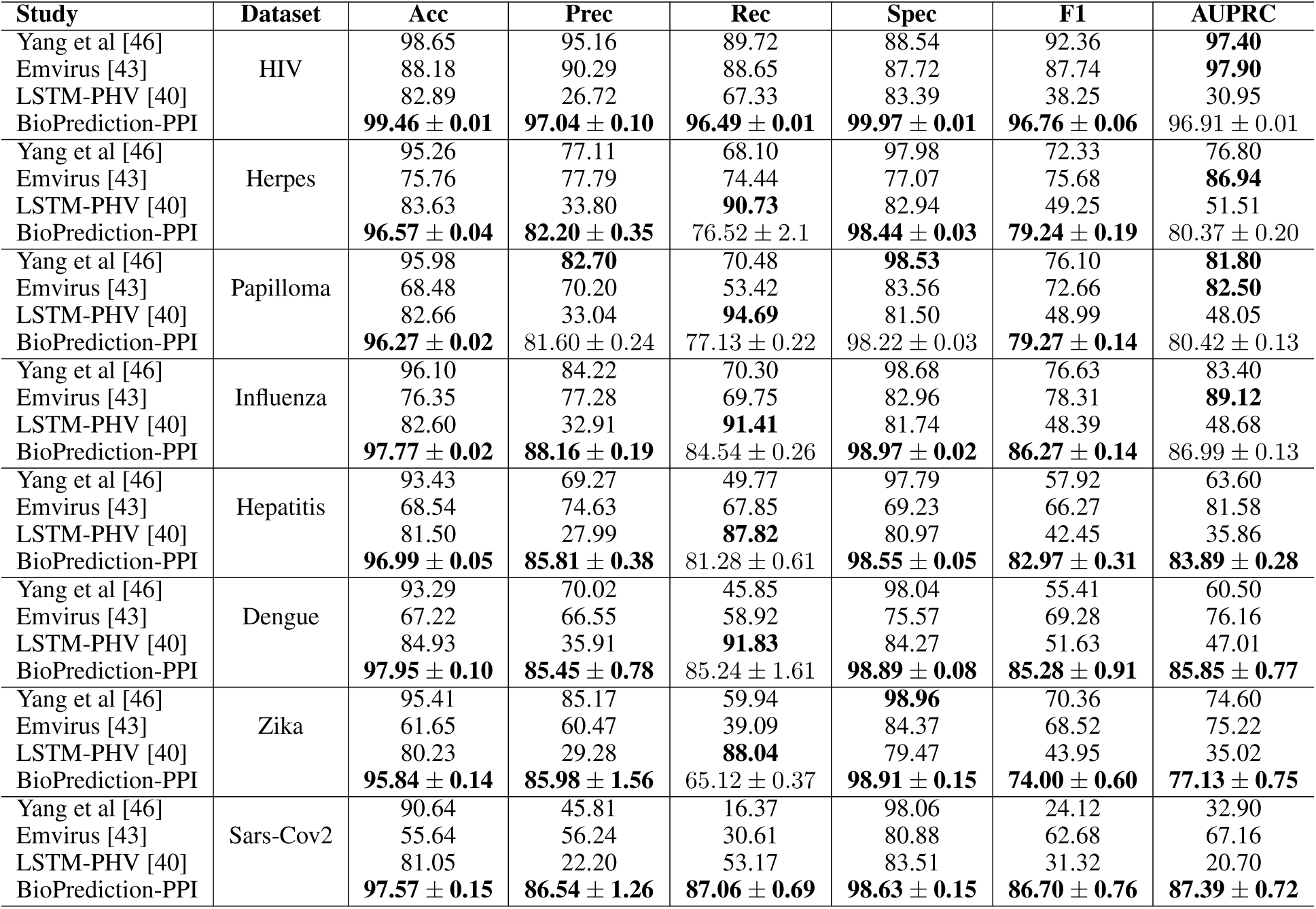
Performance in robustness experiment (HIV, herpes, papilloma, influenza, hepatitis, dengue, Zika, and SARS-CoV-2 datasets), measuring the Accuracy (Acc), precision (Prec), recall (Rec), specificity (Spec), F1-score (F1), and Area Under the Precision-Recall Curve (AUPRC). Except for LSTM-PHV [40], where performance was calculated using predictions from its web server, the performances were taken from the original articles of the respective tool.

| Study | Dataset | Acc | Prec | Rec | Spec | F1 | AUPRC |
| --- | --- | --- | --- | --- | --- | --- | --- |
| Yang et al [46] | HIV | 98.65 | 95.16 | 89.72 | 88.54 | 92.36 | **97.40** |
| Emvirus [43] |  | 88.18 | 90.29 | 88.65 | 87.72 | 87.74 | **97.90** |
| LSTM-PHV [40] |  | 82.89 | 26.72 | 67.33 | 83.39 | 38.25 | 30.95 |
| BioPrediction-PPI |  | **99.46 ± 0.01** | **97.04 ± 0.10** | **96.49 ± 0.01** | **99.97 ± 0.01** | **96.76 ± 0.06** | 96.91 ± 0.01 |
| Yang et al [46] | Herpes | 95.26 | 77.11 | 68.10 | 97.98 | 72.33 | 76.80 |
| Emvirus [43] |  | 75.76 | 77.79 | 74.44 | 77.07 | 75.68 | **86.94** |
| LSTM-PHV [40] |  | 83.63 | 33.80 | **90.73** | 82.94 | 49.25 | 51.51 |
| BioPrediction-PPI |  | **96.57 ± 0.04** | **82.20 ± 0.35** | 76.52 ± 2.1 | **98.44 ± 0.03** | **79.24 ± 0.19** | 80.37 ± 0.20 |
| Yang et al [46] | Papilloma | 95.98 | **82.70** | 70.48 | **98.53** | 76.10 | **81.80** |
| Emvirus [43] |  | 68.48 | 70.20 | 53.42 | 83.56 | 72.66 | **82.50** |
| LSTM-PHV [40] |  | 82.66 | 33.04 | **94.69** | 81.50 | 48.99 | 48.05 |
| BioPrediction-PPI |  | **96.27 ± 0.02** | 81.60 ± 0.24 | 77.13 ± 0.22 | 98.22 ± 0.03 | **79.27 ± 0.14** | 80.42 ± 0.13 |
| Yang et al [46] | Influenza | 96.10 | 84.22 | 70.30 | 98.68 | 76.63 | 83.40 |
| Emvirus [43] |  | 76.35 | 77.28 | 69.75 | 82.96 | 78.31 | **89.12** |
| LSTM-PHV [40] |  | 82.60 | 32.91 | **91.41** | 81.74 | 48.39 | 48.68 |
| BioPrediction-PPI |  | **97.77 ± 0.02** | **88.16 ± 0.19** | 84.54 ± 0.26 | **98.97 ± 0.02** | **86.27 ± 0.14** | 86.99 ± 0.13 |
| Yang et al [46] | Hepatitis | 93.43 | 69.27 | 49.77 | 97.79 | 57.92 | 63.60 |
| Emvirus [43] |  | 68.54 | 74.63 | 67.85 | 69.23 | 66.27 | 81.58 |
| LSTM-PHV [40] |  | 81.50 | 27.99 | **87.82** | 80.97 | 42.45 | 35.86 |
| BioPrediction-PPI |  | **96.99 ± 0.05** | **85.81 ± 0.38** | 81.28 ± 0.61 | **98.55 ± 0.05** | **82.97 ± 0.31** | **83.89 ± 0.28** |
| Yang et al [46] | Dengue | 93.29 | 70.02 | 45.85 | 98.04 | 55.41 | 60.50 |
| Emvirus [43] |  | 67.22 | 66.55 | 58.92 | 75.57 | 69.28 | 76.16 |
| LSTM-PHV [40] |  | 84.93 | 35.91 | **91.83** | 84.27 | 51.63 | 47.01 |
| BioPrediction-PPI |  | **97.95 ± 0.10** | **85.45 ± 0.78** | 85.24 ± 1.61 | **98.89 ± 0.08** | **85.28 ± 0.91** | **85.85 ± 0.77** |
| Yang et al [46] | Zika | 95.41 | 85.17 | 59.94 | **98.96** | 70.36 | 74.60 |
| Emvirus [43] |  | 61.65 | 60.47 | 39.09 | 84.37 | 68.52 | 75.22 |
| LSTM-PHV [40] |  | 80.23 | 29.28 | **88.04** | 79.47 | 43.95 | 35.02 |
| BioPrediction-PPI |  | **95.84 ± 0.14** | **85.98 ± 1.56** | 65.12 ± 0.37 | **98.91 ± 0.15** | **74.00 ± 0.60** | **77.13 ± 0.75** |
| Yang et al [46] | Sars-Cov2 | 90.64 | 45.81 | 16.37 | 98.06 | 24.12 | 32.90 |
| Emvirus [43] |  | 55.64 | 56.24 | 30.61 | 80.88 | 62.68 | 67.16 |
| LSTM-PHV [40] |  | 81.05 | 22.20 | 53.17 | 83.51 | 31.32 | 20.70 |
| BioPrediction-PPI |  | **97.57 ± 0.15** | **86.54 ± 1.26** | **87.06 ± 0.69** | **98.63 ± 0.15** | **86.70 ± 0.76** | **87.39 ± 0.72** |

The web server LSTM-PHV presented the lowest performance, particularly in precision, with values of around 30%. Due to its significantly lower performance compared to the other models, it was excluded from further analyses. However, it is important to note that this web server was pre-trained on human-viral interactions and is not specifically tailored to each dataset, which contributes to its lower performance.

When analyzing the results of BioPrediction-PPI, we observe that, except for the HIV dataset (with 10,000 positive interactions), which achieved 97% precision, all other precision values fall within the 80% range. This holds across datasets, from the herpes dataset with approximately 6,000 interactions to the SARS-CoV-2 dataset with around 600 interactions.

When comparing the precision performance of BioPrediction-PPI with the highest reference performance from Yang et al. [46] and Emvirus [43], the differences across the four datasets with the highest number of positive interactions (HIV - 9,880, herpes - 5,966, papilloma - 5,099, and influenza - 3,044) are +1.88, +4.41, -1.10, and +3.94, respectively. In contrast, the differences in the four datasets with fewer interactions (hepatitis - 1,300, dengue - 927, zika - 709, SARS-CoV-2 - 568) are +11.18, +15.43, +0.81, and +30.30, respectively.

Therefore, in datasets with a higher number of interactions, the precision difference is much smaller compared to those with fewer interactions. This indicates that BioPrediction-PPI achieves superior precision performance, particularly when working with a smaller number of interactions. As the number of interactions increases, its performance tends to align with that of other tools.

When performing the same comparison for sensitivity, the differences are +6.77, +2.08, +6.65, +14.24, +13.43, +26.32, +5.18, and +56.45, respectively, from the largest to the smallest dataset. In terms of sensitivity, we observe a clear distinction between the three largest datasets and the five smaller ones, with BioPrediction-PPI significantly outperforming other approaches, especially as the dataset size decreases.

### 3.3 Generalization Experiment Results

Experiment 3 aimed to evaluate the generalization, through the performance of BioPrediction-PPI in predicting PPIs between different species, an essential aspect in understanding evolutionary conservation and functional similarities among organisms. As summarized in Table 7, BioPrediction-PPI displayed competitive performance against three of the six evaluated studies, indicating its median robustness in predicting PPI between species. However, it is important to note that the three studies that outperformed BioPrediction-PPI (D-SCRIPT [35], Topsy-Turvy [34], and TUnA [24]) are DL models specifically designed to handle the complexities of cross-species interactions, even in these particular datasets. Additionally, BioPrediction-PPI demonstrated superior performance compared to PIPR [10] and DeepPPI [13], which had previously outperformed it in the first and third experiments, respectively.

**Table 7:** Performance in generization experiment, measuring the Area Under the Precision-Recall Curve (AUPRC) and Area Under the Receiver Operating Characteristic Curve (AUROC). The performance of LSTM-PHV [40] was calculated via its web server, while the others were taken from TUnA [24].

| Study | Dataset | AUPRC | AUROC |
| --- | --- | --- | --- |
| LSTM-PHV [40] | *E. coli* | 12.0 | 62.0 |
| PIPR |  | 27.1 | 67.5 |
| DeepPPI |  | 27.1 | 68.8 |
| D-SCRIPT |  | **51.3** | **77.0** |
| Topsy-Turvy |  | **55.6** | **80.5** |
| TUnA [24] |  | **67.7** | **88.3** |
| BioPrediction-PPI |  | 41.14 | 76.95 |
| LSTM-PHV [40] | *M. musculus* | 35.0 | 67.0 |
| PIPR |  | **52.6** | 83.9 |
| DeepPPI |  | **51.8** | 81.6 |
| D-SCRIPT |  | **66.3** | **90.1** |
| Topsy-Turvy |  | **73.5** | **93.4** |
| TUnA [24] |  | **88.5** | **97.8** |
| BioPrediction-PPI |  | 46.49 | 83.93 |
| LSTM-PHV [40] | *D. melanogaster* | 14.0 | 62.0 |
| PIPR |  | 27.8 | 72.8 |
| DeepPPI |  | 23.1 | 65.9 |
| D-SCRIPT |  | **60.5** | **89.0** |
| Topsy-Turvy |  | **71.3** | **92.1** |
| TUnA [24] |  | **85.5** | **97.1** |
| BioPrediction-PPI |  | 35.82 | 79.85 |
| LSTM-PHV [40] | *C. elegans* | 17.0 | 67.0 |
| PIPR |  | 34.6 | 75.7 |
| DeepPPI |  | 25.2 | 67.1 |
| D-SCRIPT |  | **55.0** | **85.3** |
| Topsy-Turvy |  | **70.0** | **90.6** |
| TUnA [24] |  | **83.4** | **96.0** |
| BioPrediction-PPI |  | 39.22 | 81.31 |
| LSTM-PHV [40] | *S. cerevisiae* | 14.0 | 64.0 |
| PIPR |  | 23.0 | 71.8 |
| DeepPPI |  | 20.1 | 65.2 |
| D-SCRIPT |  | **39.9** | **79.0** |
| Topsy-Turvy |  | **53.4** | **85.0** |
| TUnA [24] |  | **64.1** | **89.8** |
| BioPrediction-PPI |  | 28.55 | 76.53 |

These results point out potential directions for future improvements of our framework in cross-species scenarios. Therefore, this experiment provides initial evidence that BioPrediction-PPI may outperform studies not explicitly designed for this subtask, specifically those involving PPI prediction. Moreover, the results underscore the complexity of cross-species PPI prediction, where evolutionary distances and species-specific interaction dynamics can significantly impact the predictive accuracy.

## 4 Interpretability Experiment Results - Case Study

This section discusses the interpretability, so we present a use case that leverages a viral-human interaction dataset, specifically the Influenza dataset, to showcase the versatility of the developed framework. The performance results are presented in Figure 5. Bioprediction-PPI performance was marginally lower than in Table 6; this is likely because it was trained on only 6 out of 10 of the data.

**Figure 5:**
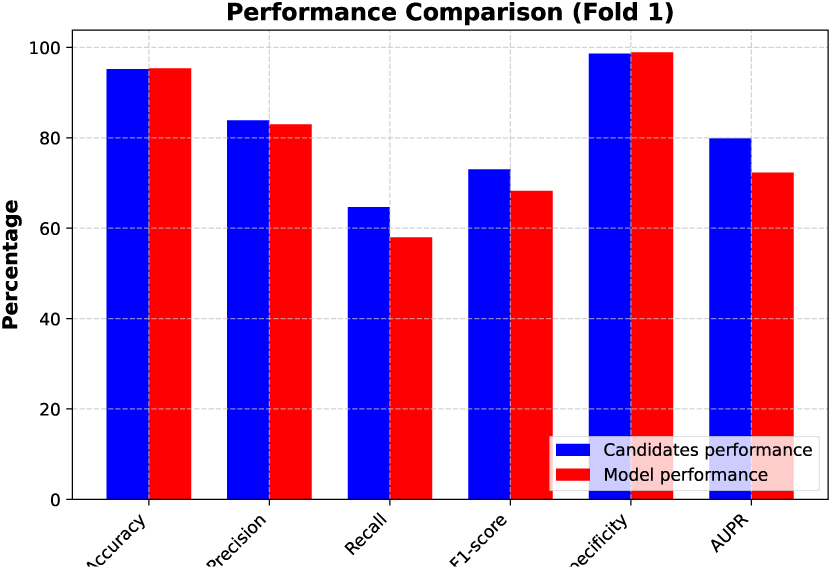
Performance of the model constructed with 6/10 of the Influenza dataset and the others 4/10 of the data used as candidate interaction pairs.

**Figure 6:**
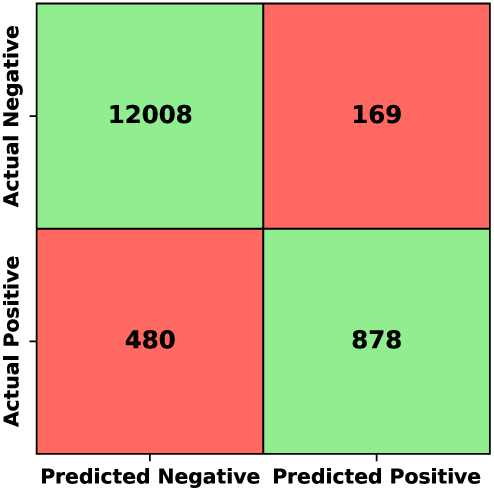
Confusion matrix of predictions for candidate interaction data in the case of study experiment.

The slightly superior performance observed in the candidate interactions compared to the model’s estimated performance is due to the imposed constraint, which requires candidate interactions to involve proteins present in the training set. As a result, proteins that form only one interaction pair are included in the overall training subset and later also in the test set used to estimate the model’s performance. However, since the model has limited information about these proteins, its performance in these cases is lower, which ultimately reduces the overall performance.

At this stage, with all the predictions of candidate interactions (using a threshold of 0.5), the user received 1,047 new positive interactions, 878 true and 169 false. Thus, approximately 4 out of every 5 interactions were correct, as measured by the approximately 83% precision. This shows that, by testing just 1,047 pairs in the laboratory, the user achieves a 52% increase in the number of known positive interactions. This is better than expected, considering that in the model’s training set, there were only 1686 positive pairs, and now the user discovered new 878 positive interactions. The corresponding confusion matrix, illustrated in Table 6, details the correct and misclassifications.

### 4.1 Complementary Graphs

The next results represent what a user would obtain when using BioPrediction-PPI, based on the measured metrics. However, since other models also demonstrate competitive performance compared to the developed framework, these outcomes could similarly be achieved using other previously presented tools. In this context, we will analyze the additional outputs provided by BioPrediction-PPI to see what other insights the framework offers that could assist users in decision-making. These outputs are visual reports that explain the behavior of the trained model, always based on the data used for modeling. A detailed description of these visualizations can be found in the BioPrediction-PPI Reports subsection. A first graph is the **distribution of predicted scores** by the model for each class generated based on the test set, which is represented in Figure 7.

**Figure 7:**
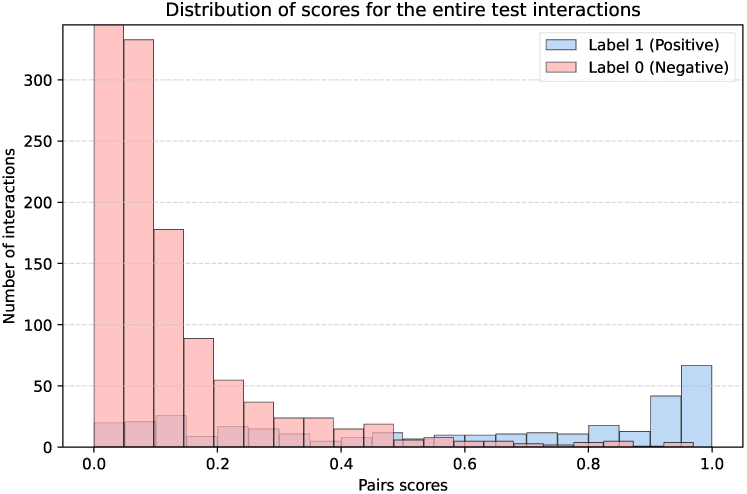
Distribution of the predicted scores by the model in the test set, separated by class. The Y-axis of the distribution is limited to 250 interactions, as the first bar represents approximately 3,000 interactions, thereby restricting the visibility of other parts of the graph.

We observed that both the positive class distribution and the negative class distribution exhibit a long-tailed and unilateral pattern. The positive distribution has its peak near the score of 1.0, with a tail extending towards the score of 0.0, while the negative distribution follows the opposite behavior. This pattern suggests that the model concentrates its positive predictions at higher scores, confidently identifying the most likely interacting protein pairs in this region.

However, it still assigns positive scores throughout the entire space, reflecting cases with higher uncertainty or where some interactions do not perfectly follow the learned behavior.

Moreover, it is noted that the two distributions intersect, indicating that there are protein pairs from both classes with the same score, although in different quantities. However, at the extremes, these distributions show the least intersection, making them the regions of highest precision for the model. Finally, this graph aims to illustrate the behavior of the predictive model, which assigns a score between 0 and 1 to each protein pair.

#### 4.1.1 Precision Graphs

Next, we analyze the model’s metrics more deeply, starting with the precision of the positive class, which is the percentage of correct predictions when the model classifies a pair as positive. The **precision by threshold interval** graph, described earlier in the methodology, separates the model’s predictions into different precision regions. In Figure 8a, we present the graph generated by BioPrediction-PPI during modeling. Since we know the labels of the candidates in this experiment, a graph for these data was also constructed (see Figure 8b).

**Figure 8:**
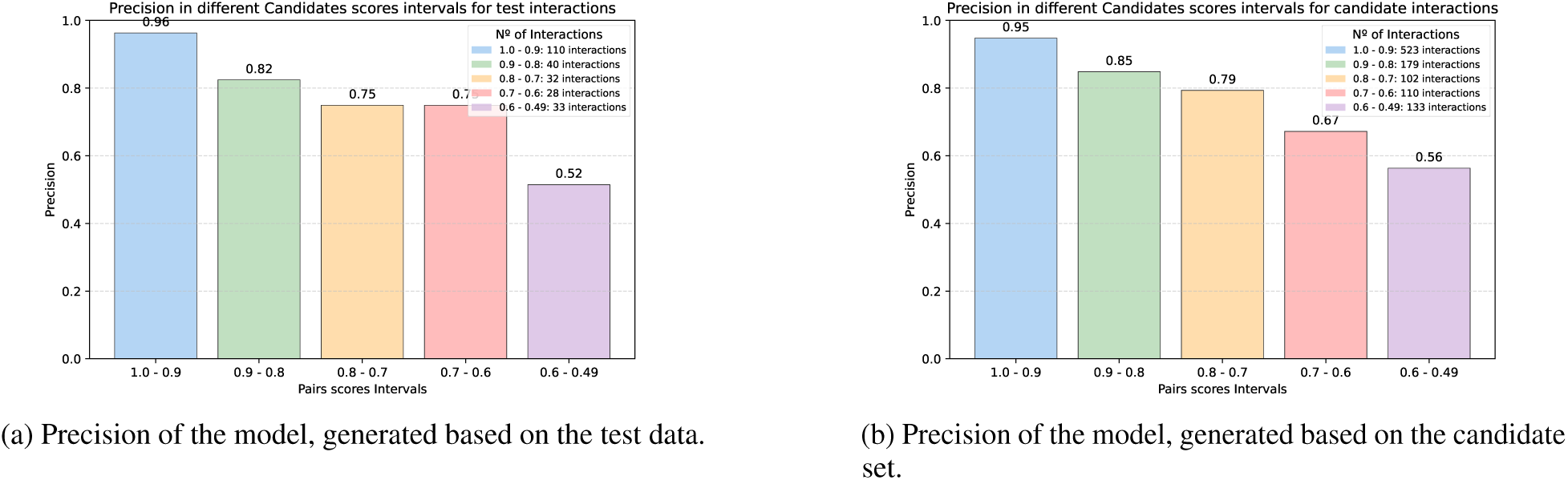
Precision of the model’s positive class by predicted score interval, in the test (used for final performance estimation) and candidate set.

First, in Figure 8a, we observe that the largest subgroup of pairs predicted as positive by the adjusted model has scores between 1.0 and 0.9, corresponding to approximately 110 predictions (or 45% of the predicted positive interactions). In this range, precision reaches 96%, a value higher than the model’s overall average on the test set, which is 83%. This indicates that the model identified this subset of predictions with high confidence, making these pairs strong candidates for experimental testing.

In the next interval (0.8–0.9), precision remains high, reaching 82%, which is close to the estimated average. On the other hand, the next three intervals (0.5–0.6, 0.6–0.7 and 0.7–0.8), which account for 38% of the interactions predicted as positive, show a significant drop in precision, ranging between 75% and 52%. This suggests that predictions within this range may not be the most reliable for experimental validation, as the precision is relatively low. Overall, when considering all predictions in the range from 1.0 to 0.5, the average precision is 83%. However, this analysis allows us to understand the precision profile across different thresholds, which can contribute to a better understanding of the model’s general behavior before its application.

The analysis of Figure 8b, which represents the predictions in the candidate interaction set, shows that the patterns identified in Figure 8a remain consistent. The highest number of pairs and the highest precision are still concentrated in the 1.0 to 0.9 range. This is followed by an interval with precision close to the estimated average, while in the remaining intervals, precision progressively decreases to approximately 50%.

This result suggests the reliability of the experimental planning based on Figure 8a, when applied to new interactions of the same nature as those used in the modeling. Furthermore, it enables the visualization of the distribution of predictions, allowing for the strategic selection of the most promising pairs for experimental testing. E.g., by only selecting interactions with the highest precision, better resource management can be achieved.

Complementary to the precision analysis, we present the **precision by threshold** graphs, constructed as described in the methodology. These graphs show the precision considering all predictions from 1.0 down to a specific score threshold. By observing Figure 9a, we can see that in the range of 1.0 to 0.9, the precision is approximately 96%, which corresponds to the value estimated in Figure 8a, as expected. When we expand the analysis to the range of 1.0 to 0.8, the precision remains close to 93%. This result occurs because this new range predominantly includes interactions from the first range, as there are 110 interactions in the 1.0-0.9 range and only 40 interactions in the 0.9-0.8 range. Therefore, when the precision is calculated including both ranges, the value is closer to the first range.

**Figure 9:**
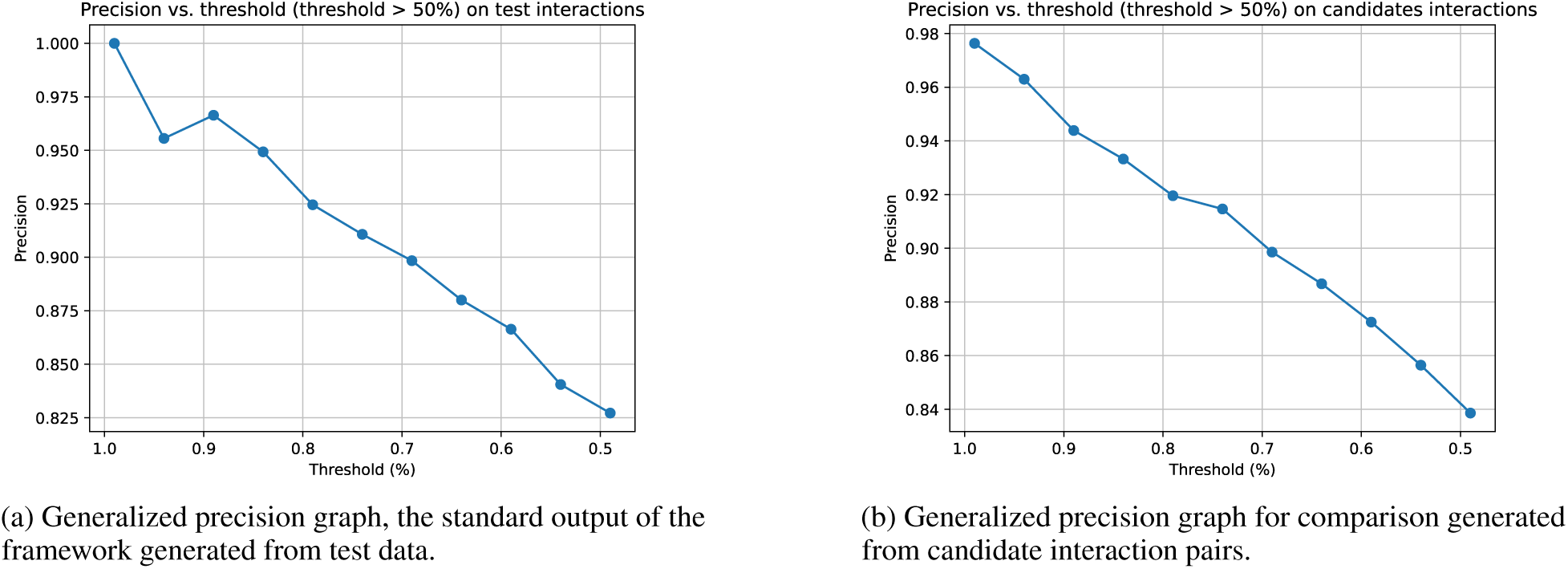
Generalized precision graph, representing precision relative to the threshold, in the test data (used for final performance estimation) and candidates interactions.

Moving forward, when the next interval (0.8–0.7) is added, the overall precision drops to 90%, although this interval has an isolated precision of 75%. This relatively modest reduction occurs again due to the smaller number of interactions in this interval compared to the previous ones, as there are only 32 interactions in this range. As the last intervals are included, the overall precision gradually approaches the model’s average. This graph allows us to visualize how precision changes as new intervals with lower performance are added. In the analyzed case, the reduction in precision from the 0.8 to 0.7 range is only 3%, which could be an acceptable trade-off, especially if it favors the discovery of new positive interactions.

Meanwhile, when analyzing the graph generated from the candidate interactions in Figure 9b, we observe that the behavior remains similar to the one seen in the graph based on the test set. This result reinforces the reliability of the graphs generated from the test set to estimate the model’s performance on new interactions. In summary, while Figure 8a highlights the regions of highest precision of the model, Figure 9a enables the evaluation of the impact of including different thresholds on the precision within a given range. Thus, these analyses help define the most appropriate cutoff point for each application, enabling more strategic and efficient experimental planning and highlighting an advantage of BioPrediction-PPI over other tools with similar performance.

#### 4.1.2 Recall Graph

Another metric that can be studied is sensitivity, which is the percentage of actual positive interactions correctly classified as positive. The **sensitivity by threshold** graph is described in the methodology. However, given the construction of the graph defined by Figures 10a and 10b, we observe that before the red line, all pairs are predicted to be positive. At this stage, the curve increases rapidly, indicating a high rate of true positives relative to the percentage of the dataset. However, after the red line, the rate of true positives per percentage of the dataset begins to progressively decrease. This suggests that to increase the true positive rate, it becomes necessary to test more and more data. Based on an analysis of its resources, the user can determine the extent to which working in the saturated regime is useful for obtaining a minimum percentage of the total positive interactions.

**Figure 10:**
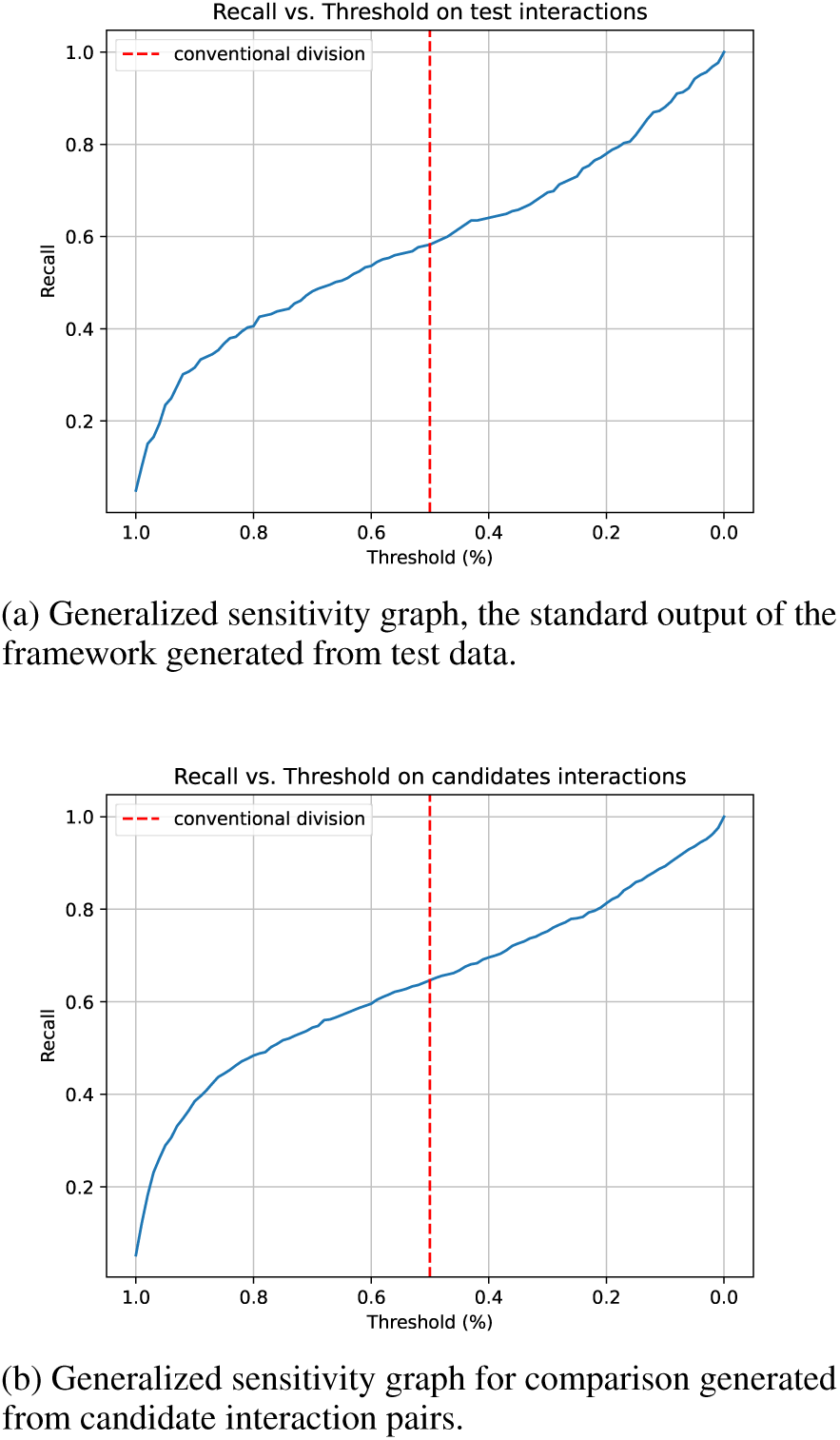
Generalized sensitivity graph representing sensitivity relative to the threshold in the test data (used for final performance estimation), and in the candidates’ interactions.

To support this, analysis of the candidate interaction data visualized in the graph (Figure 10b) reveals a behavior similar to that of the test data graph (Figure 10a). This similarity strengthens the case for using the test-generated graph as an estimate for predicting new data. The graph 10a generated during the BioPrediction-PPI validation allows one to determine that, to obtain approximately 80% of the positive interactions, it is necessary to test until the threshold score of 0.2, which implies considering all predicted interactions as positive and a little more. Therefore, BioPrediction-PPI provides a visual means to understand the model’s sensitivity, enabling users to plan the size of their experiment more efficiently.

This type of information can be relevant if the user wants to obtain a measure of the completeness of the interactions captured, considering a given threshold. For example, it is possible to understand that, when using 0.9 as the threshold, approximately 30% of all interactions will be included (and, as seen earlier, with about 96% precision). Moreover, these graphs enable users to determine the threshold at which they should test to achieve a certain percentage of interactions in their experiment. This can be useful to ensure a minimum fidelity of the interaction network under study, ensuring that a sufficient number of interactions are considered for subsequent analyses.

#### 4.1.3 Biological Insights Graphs

The interpretability report employs various graphs to facilitate the understanding of the impact each feature has on the model’s decision-making process. Figures 11a and 11b illustrate this approach, with one graph focusing on the final classification and another representing a partial model.

**Figure 11:**
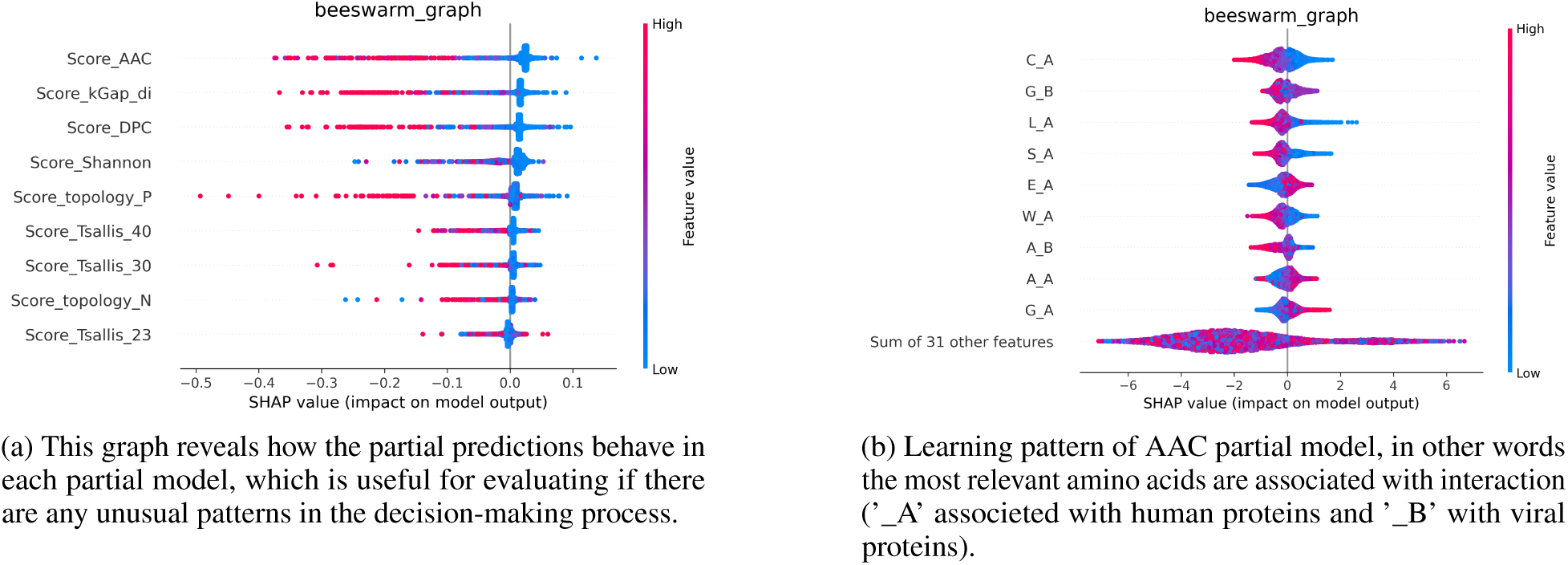
Interpretability output graphs from BioPrediction-PPI.

In the graph, each sample is represented by a color gradient ranging from red to blue, where red signifies a high value for the feature in question and blue indicates a low value. The horizontal distance of each point from the center of the distribution (0.0) shows how strongly the feature contributed, either positively (positive SHAP values) or negatively (negative SHAP values), to the final classification for a specific class. On the left side of the graph, the nine most impactful features are listed, along with an individual analysis of each.

In Figure 11a, we analyze the final model, which combines the decisions of the partial models. Each partial model predicts protein pairs with a score between 0 and 1, where values greater than 0.5 indicate interactions between proteins, while lower values indicate the absence of interaction. Among the partial models, the most relevant one, according to the figure, is the one based on AAC (Amino Acid Composition) features, which counts the frequency of each amino acid in both proteins.

Observing the AAC model, we see that in the final decision, all high predicted values from this model are considered to contribute to the prediction of the positive class, demonstrating the consistency of the initial positive interaction predictions made by this model. However, the model also identifies a range of low values, represented in blue, that contribute to the prediction of the positive class. This suggests that these pairs were initially misclassified, but during the final decision, the new model attempted to correct these incorrect decisions.

Unlike the AAC model, the model based on Tsallis features shows high values that contribute to both the positive and negative class predictions, indicating less consistency in its initial predictions. In other words, when analyzed in isolation, the Tsallis model is less consistent than the AAC model. In Figure 11b, we observe the more general patterns learned by the AAC-based partial model. The label ’_A’ is associated with human proteins, as in the interaction table used for training, human proteins are listed in the first column. Consequently, the label ’_B’ is associated with viral proteins, as they correspond to the second column in the training table.

When analyzing the three most frequent amino acids in human proteins as identified by the model, we find cysteine (C), serine (S), and leucine (L). The first two have negatively polar side chains without charge, while the latter is a bulky, nonpolar amino acid. The high frequency of these amino acids in the primary structure is more associated with non-interaction between proteins. Among the most relevant features, only two amino acids from the viral structure stand out: alanine (A) and glycine (G). Both are nonpolar amino acids with small side chains, and their high frequency is also more associated with non-interaction. However, when analyzing human proteins, we observe that elevated frequencies of A and G are associated with interactions.

This type of plot is valuable for understanding what the model has learned, revealing data patterns that can direct the user’s attention to specific regions of the proteins, where certain characteristics may be more relevant for interaction or non-interaction. A similar analysis can be performed with the DPC (Dipeptide Composition) partial model, which counts the frequency of each amino acid pair in the primary structure of the proteins, potentially offering even more insights into protein interaction patterns.

A key point is that users have the option to rely exclusively on a single partial model, using its decisions to guide their experiments, even if this may lead to a slight reduction in performance. This approach simplifies the decision-making process and enhances interpretability. However, it must be mentioned that not all features have a straightforward biological significance. Thus, a critical consideration for future iterations is to prioritize the development of features with greater biological relevance, potentially replacing those that are currently less interpretable.

### 4.2 A Protein Analysis

In this subsection, we performed a manual analysis of predictions for new, unlabeled interactions for a single protein, selecting the protein with a high number of positive interactions in the candidate interaction dataset. The chosen protein is an Influenza virus protein identified by the code P03496. First, we will visualize, in Figure 12a, the distribution of the predicted scores by the model for the positive and negative classes of the pairs involving this protein, generated based on the test set.

**Figure 12:**
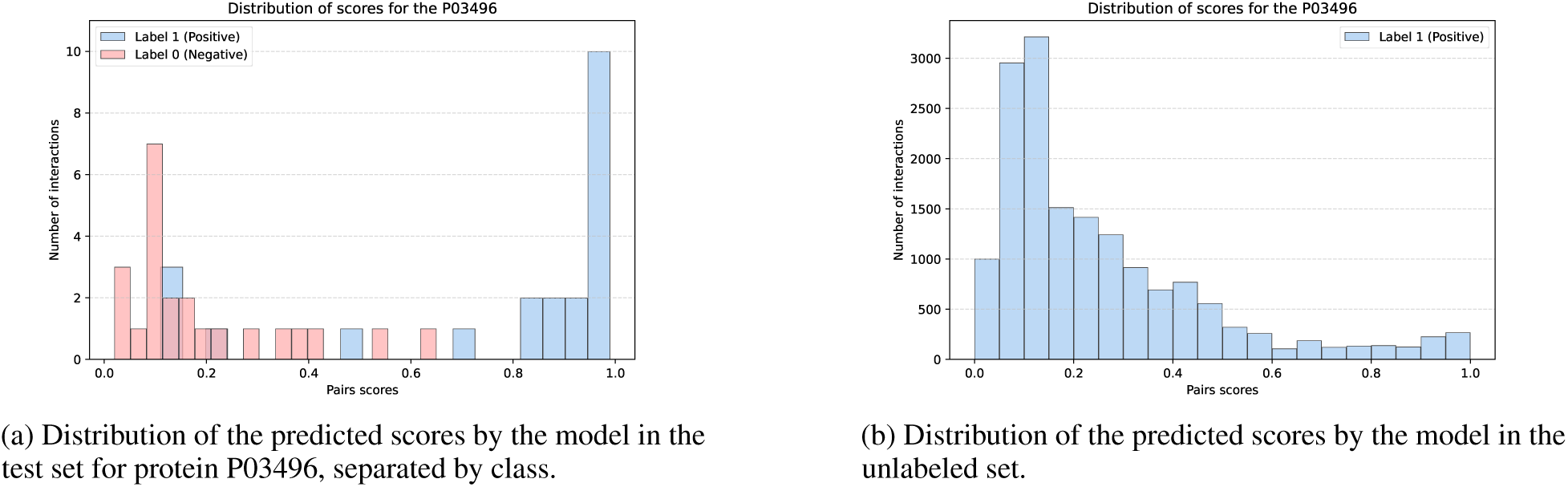
Distribution of the predicted scores for protein P03496.

In Figure 12a, despite the limited number of interactions to generate the distribution, we observe that it follows the characteristic behavior of long-tail and skewed distributions, similar to what is observed for the model as a whole in Figure 7.

Next, for the same protein, we selected all unlabeled pairs between this viral protein and human proteins in the original dataset, taken from the study by Yang et al. [46], resulting in 16,190 new unlabeled pairs. We then used the first fold of the predictive model again to predict these unlabeled pairs. Subsequently, we recalculated the distribution of predicted scores. However, since these pairs are unlabeled, no separation by class is available, as can be seen in Figure 12b.

In Figure 12b, we can assume that the distribution is the sum of the long-tail, unilateral distributions of the positive and negative classes, similar to what would be generated by summing the distributions observed in Figure 7. This behavior in Figure 12b is as expected, as, in addition to the shape, there is a larger number of interactions labeled as negative, given that a protein typically interacts with only a fraction of the existing proteins. Based on the entire analysis of the model in question, we can conclude that the model’s metrics are valid. That is, of the 1,832 interactions predicted as positive (those with a score above 0.5), 80% should be correct. Additionally, considering a threshold score of 0.9, we have 497 protein pairs, of which we expect approximately 95% to be correct.

Seeking validation for these new predictions, the originally unlabeled interactions in the Yang et al. [46] dataset were searched in the IntAct interaction database [11]. This search resulted in 196 of these pairs being found, of which 174 were classified as positive (89% sensitivity), with 136 of these classifications having a score above 0.9. This reinforces that the majority of predictions occur in this region, which is also the most precise.

## 5 Discussion

First, several aspects of the framework developed in this study were investigated, with these analyses focusing on its automation, robustness, generality, and interpretability. In the first experiment, we evaluated the performance of the BioPrediction-PPI model, an end-to-end framework, against three different benchmarks. The first benchmark consisted of 25 distinct studies, while the other two benchmarks comprised 12 studies each. After conducting the experiments, the results indicated that, overall, BioPrediction-PPI was competitive, except in a few specific cases.

The only models that outperformed BioPrediction-PPI were: the PIPR model [10], trained on the yeast dataset; the LBG+BBA model [52], trained on the H. pylori dataset; and the DeepFE-PPI [47] and DeepPPI [13] models when evaluated on the human benchmark. In these cases where the competing models showed superior performance, the difference was relatively small, for example, with their accuracy being only 2% to 4% higher than that of BioPrediction-PPI.

Of these four studies, only PIPR [10] also adopts an automated approach 1. However, this is a DL model in both the feature extraction and prediction stages, making it inherently less interpretable. Alternatively, although PIPR [10] is automated for training, it lacks an automated component that enables the adjusted model to be used for predicting new interactions, and it does not provide supplementary reports to users to clarify the decision-making process or offer guidance on how to utilize the model.

Also, these models did not exhibit superior performance across the other datasets. For example, in the generalization experiment, BioPrediction-PPI did not perform worse than PIPR [10]. Furthermore, on the yeast dataset, BioPrediction-PPI again did not show inferior performance compared to the LBG+BBA [52], DeepFE-PPI [47], and DeepPPI [13] models. Overall, all models compared to BioPrediction-PPI showed equal or inferior performance on at least one of the tested datasets.

These results demonstrate that it is possible to employ an automated model that remains competitive and, in many cases, outperforms other models. Notably, no DL approaches were used in this pipeline. This decision was driven by the goal of preserving interpretability, an essential aspect of understanding and applying the model in practical contexts. Moreover, the results suggest that we possess a solid understanding of the PPI prediction problem and a strong modeling capability. This knowledge allowed us to select a specific pipeline that proved to be as effective as several alternative architectures with comparable performance, despite being based on different feature extraction strategies and predictive models.

In the second experiment, we evaluated the robustness of the models using various datasets of human–viral interactions, which presented imbalanced conditions and variability in dataset size. BioPrediction-PPI demonstrated equal or superior performance compared to all models from previous studies that addressed these datasets, with notable effectiveness on datasets with fewer interactions, where the model proved particularly capable of learning patterns from a limited amount of data.

This performance difference is especially significant in the SARS-CoV-2 dataset (containing 568 positive interactions), where the Emvirus study [43] achieved a precision of 56.24% and a recall of 30.61%. In contrast, BioPrediction-PPI achieved 86.54% precision and 87.06% recall, demonstrating vastly superior performance compared to the results reported by other studies. This specific case represents an extreme scenario, with a performance gap of 30% or more, highlighting the model’s ability to handle small-scale datasets effectively. However, as the datasets increase in size, the performance of the models tends to converge. Even so, the results highlight the robustness of the developed framework, which is capable of effectively learning across different scenarios, taking into account both dataset size and class imbalance.

This can be explained by the fact that the model utilizes features based on both the dataset’s topology and the molecular structure. For smaller datasets, there may be insufficient data for training, making it more challenging to infer interaction patterns solely based on primary protein structures, such as identifying important amino acids, since these patterns are complex. However, the interaction network, despite its smaller size, can provide a solid characterization of the problem, allowing the model to learn patterns with limited data. This enables the model to perform well even in data-scarce scenarios by leveraging topological features to support its predictions.

In the third experiment, focused on generalization, we evaluated the ability of the trained models to make predictions in previously unseen contexts. In this setting, BioPrediction-PPI proved to be competitive, outperforming 3 out of the 6 models. Therefore, it does not appear to be the best option for generating models with high generalization capacity.

Instead, its application is more suitable for datasets related to the proteins of interest, particularly when related or homologous proteins are present.

In this context, applying models specifically trained to handle a given set of proteins offers several advantages. The main one is that the performance estimated during model training tends to closely reflect the actual performance when the model is applied to predict new interactions related to the same set or target species. In contrast, a model designed to be general may face challenges in estimating its expected performance on a new set of proteins or a different target species.

For example, the Tuna model [24], which shows good generalization ability, achieved an AUPRC of 88.5 for the species *M. musculus* and 64.1 for *S. cerevisiae*, resulting in a relatively wide range between the best- and worst-case performance. Given that users may be working with proteins from any species, there is no clear method to estimate the final performance of such a general model. On the other hand, a model specifically tailored to a given context can have its performance optimized for that particular subproblem. This lack of interpretability regarding how performance may vary in new contexts represents a significant limitation of general-purpose models.

Still within the context, the fourth experiment aimed to explore the interpretability of the framework by analyzing the dataset of protein interactions between humans and the influenza virus. During this analysis, it became evident that the model’s estimated performance, traditionally evaluated using a 0.5 threshold, follows a particular distribution that can be further explored due to the model’s interpretability. Therefore, between two models with no significant differences in performance, the one that allows for a deeper exploration of this distribution can provide greater support for experimental planning.

Consequently, we explored the model’s precision as shown in Figure 8a, revealing its distribution. Instead of assuming that the estimated performance is uniformly distributed, that is, assuming a constant precision across the 0.5 to 1.0 interval, we measured the precision across several subintervals, identifying where it is above or below average. In the specific case studied, the average precision was 83%, but in the 0.9 to 1.0 range, nearly half of the predictions were concentrated, resulting in a precision of 96%. This indicates a much higher reliability in that region, suggesting it as a preferential range for selecting pairs for new experiments. Conversely, in the 0.5 to 0.8 range, approximately one-third of the protein pairs were classified with a precision between 75% and 52%. Providing users with access to this type of information enables them to make informed decisions about whether to include these pairs in subsequent experiments.

Moreover, any decision to adopt more or fewer interactions predicted within the 1.0 to 0.5 interval affects the model’s recall. In this sense, a graph is also provided that explores the model’s sensitivity, offering an estimate of this metric for each threshold interval, as shown in Figure 10a. This type of information allows for finer control over the model’s decision-making process. Additionally, a set of graphs related to feature importance is available, which, combined with the fact that some features have clear biological meanings, can help generate hypotheses about the interactions being studied.

Finally, predictions for a single protein were evaluated, and it was found that they follow a pattern similar to that of the model as a whole, as shown in Figure 12a. The positive proteins were predicted with scores that follow a distribution centered around 1.0, with a tail extending towards the value 0. This indicates that, for this protein, certain interaction patterns were learned, classifying these interactions with values close to 1, while almost no non-interacting pairs received this score.

Additionally, there are less clear interaction patterns for the model, such as values between 0.8 and 0.5, where the model still predominantly classifies interactions as positive. Still, a considerable number of negative interactions occur within this range. Finally, the model also classified some interactions as negative, suggesting that it did not learn or effectively identify certain interaction patterns.

When the prediction of all unlabeled human protein pairs and those of this virus is made within the original dataset, as shown in Figure 12b, the distribution of scores follows a pattern that can be interpreted as the estimated distribution in Figure 12a, but with both positive and negative classes combined, in a real-world scenario where there are far more negative interactions than positive ones.

We can say that these predictions align with the behavior initially estimated. Upon searching for these unlabeled interactions in other databases, about 200 were found, 89% of which were predicted as positive, with 82% of the predicted positive interactions falling within the 1.0 to 0.9 score range, a region where interpretability previously indicated that the model concentrates the highest number of positive interactions with the highest precision. Therefore, this example illustrates This example shows the interpretability facility provided by BioPrediction-PPI and suggests a potential practical application scenario for the framework, such as discovering hidden interactions within a known interaction network.

Additionally, unlike competing models, BioPrediction-PPI offers an automated solution for both building predictive models and predicting new protein pairs, eliminating the need for manual interventions or programming knowledge, and improving predictive performance. This feature makes the framework more accessible to researchers without ML expertise, expanding its potential for application. Another distinguishing feature of BioPrediction-PPI is the incorporation of interpretability-focused graphs, which enable a better understanding of the model’s decisions and facilitate experimental planning. The utility of these graphs was thoroughly explored in the interpretability experiment, demonstrating their relevance for result analysis.

Throughout the study, only datasets that had already been used in other studies were employed to ensure that at least one performance reference was always provided, thereby avoiding the presentation of decontextualized results. For example, although it is possible to achieve 99% performance on certain datasets, other models may also achieve similar results. Therefore, the study focused exclusively on datasets that already had performance references from other studies, ensuring comparisons with established benchmarks. Therefore, this study did not address the construction of datasets, which involves research questions related to understanding what aspects a dataset needs to have to teach an ML model to identify interaction patterns of one or more target proteins.

Thus, if a user has a target protein, what types of interactions should they seek in databases to create an optimal model that can be applied to their protein? Various aspects can be considered in this construction, such as the function of the interacting proteins and their similarity to the target protein. In the datasets used here, these aspects were not incorporated into the formulation of the datasets, as they are samples taken from subproblems. No dataset was specifically designed to predict a group of previously unlabeled target interactions. This, naturally, opens the door for future work that investigates the formulation of datasets for molecular target inference, as this study showed that, given a dataset, modeling can be carried out automatically, without performance loss, when compared with models created by ML experts.

## 6 Conclusion

This study experimentally evaluated the performance, robustness, generalization, and interpretability of an automated tool for end-to-end PPI, BioPrediction-PPI. The results show that it is possible to develop an efficient framework that performs equal to or better than available alternatives across various datasets, such as human, yeast, bacteria, and viruses. Given a set of positive and negative interactions as input, BioPrediction-PPI has been shown to perform all intermediate processes required for the development of a predictive model. This allows users to employ it as a framework for fitting customized models by providing datasets that are representative of their specific problems. An important finding is that the framework delivers competitive performance without relying on DL techniques for feature extraction or model development. Furthermore, BioPrediction-PPI performs well in eukaryotic species (human data), in prokaryotic cases (*H. pylori* data), as well as in viral interactions and provides performance graphs to help users apply the model to their specific problems. These graphs estimate the behavior of precision and recall at different thresholds and can be used to identify signs of underfitting or overfitting. They also allow for the selection of a subset of predictions to prioritize either precision or recall. This makes them a versatile tool for scenarios that require high precision, as well as those where high recall is essential to identify as many positive interactions as possible. Thus, BioPrediction-PPI can assist biologists and researchers who are not computer experts in the development of AI models. Its accessibility and effectiveness offer the potential to significantly advance the exploration and understanding of PPIs, potentially accelerating research and development in problems related to protein interactions, such as drug discovery, disease mechanism studies, and therapeutic target identification.

## 7 Data Availability

Our framework is available for download on the following website: https://autoaipandemics.icmc.usp.br/. A preliminary version of this study was selected as the global winner of the Undergraduate Awards in 2024, with its earlier version available at: https://gua.soutron.net/Portal/Default/en-GB/RecordView/Index/3202.

## 9 Author Contributions Statement

**Bruno Rafael Florentino:** Conceptualization, Data curation, Formal analysis, Methodology, Software, Validation, Visualization, Writing – original draft, Writing – review & editing. **Robson Parmezan Bonidia:** Conceptualization, Formal analysis, Funding acquisition, Methodology, Project administration, Supervision, Validation, Writing – original draft, Writing – review & editing. **Ulisses Rocha:** Formal analysis, Validation, Writing – original draft, Writing – review & editing. **André C.P.L.F. de Carvalho:** Conceptualization, Formal analysis, Funding acquisition, Project administration, Supervision, Writing – original draft, Writing – review & editing.

## 10 Author Biographies

Bruno R. Florentino graduated in physical and biomolecular sciences at the São Carlos Institute of Physics (IFSC) University of São Paulo (USP). He is currently pursuing a Ph.D. degree in computer science and computational mathematics at the University of São Paulo - USP. His main research interests are in Machine Learning, Bioinformatics, and Molecular Biology.

Robson P. Bonidia received a Ph.D. degree in computer science and computational mathematics from the University of São Paulo - USP. Currently, he is a Professor at the Federal University of Technology-Paraná (UTFPR). His main research topics are computational biology and pattern recognition, feature extraction and selection, metaheuristics, and AI for the social good.

Ulisses N. da Rocha received his Ph.D. in Microbial Ecology from the University of Groningen (The Netherlands) in 2010. In 2017, he joined the Department of Environmental Microbiology at the Helmholtz Centre for Environmental Research - UFZ (Leipzig, Germany) as a junior group leader. In 2021, he became a senior scientist at UFZ. In addition, he has led the Microbial Data Science group at the UFZ since 2017 where he works at the intersection of Microbial Ecology, Bioinformatics, and Computational Biology.

André C.P.L.F. de Carvalho is a full professor at the Department of Computer Science, University of São Paulo. He is the Dean of the Mathematics and Computer Science Institute of the University of São Paulo, ICMC-USP. His research interests are in machine learning, data mining, and data science.

## 12 Key Highlights

- Proposal of a new framework, BioPrediction-PPI, that expands the use of predictive models for protein-protein interactions (PPIs), by researchers who are not ML-experts.
- BioPrediction-PPI, despite being an automated approach, achieves competitive performance when compared with manually designed, complex architectures, highlighting its robustness and practical applicability.
- Providing evidence that PPI prediction may depend more on the characteristics of the input datasets than on the continuous redesign of predictive methods, suggesting that standardized and well-understood pipelines can be effective across different PPI contexts.
- BioPrediction-PPI is publicly available, fostering reproducibility and accessibility for the scientific community.

## Footnotes

1 https://autoaipandemics.icmc.usp.br/

2 https://github.com/0nurB/BioPrediction-PPI

